# Basophilic Erythroblast Emerges as the Key Turning Point in Polycythemia Vera

**DOI:** 10.64898/2026.09.01.748468

**Authors:** Deniz Konak, Pınar Pir

**Affiliations:** Gebze Technical University, Department of Bioengineering, Kocaeli, Türkiye

**Keywords:** Polycythemia vera, JAK2^V617F^, basophilic erythroblast, erythropoiesis, single-cell RNA sequencing, myeloproliferative neoplasm

## Abstract

Polycythemia vera (PV) is a rare, chronic myeloproliferative neoplasm driven by the JAK2^V617F^ mutation and characterized by uncontrolled erythroid proliferation. Although the mutation arises in hematopoietic stem cells, the differentiation stage at which its transcriptional consequences first become biologically meaningful has remained undefined. Using a multi-layer transcriptomics integration approach that combined differential gene expression, NicheNet ligand-receptor analysis, pseudotime trajectory inference, and CNV profiling on scRNA-seq data, alongside bulk transcriptome validation, we identified basophilic erythroblasts as the critical transition point at which JAK2^V617F^ shifts from a genomically present but transcriptionally silent state to an actively trajectory-altering and treatment-responsive disease driver. Differential expression revealed a qualitatively distinct disease signature at this stage, including ERFE-mediated iron dysregulation, MAP2K2-driven RAS/MAPK co-activation, and epigenetic reprogramming. NicheNet showed the establishment of a TGFβ superfamily and chemokine-driven niche-remodeling axis, and pseudotime analysis demonstrated that basophilic erythroblasts are the first erythroid population to exhibit condition-dependent trajectory divergence, whereas earlier progenitors showed none despite carrying the mutation. Interferon-α treatment showed its broadest counterresponse at this stage but declined sharply thereafter, identifying basophilic erythroblasts as both the principal therapeutic target and the point of maximum vulnerability in PV.

## Introduction

Polycythemia vera is a chronic myeloproliferative neoplasm driven by the JAK2^V617F^ mutation, which causes constitutive activation of the JAK-STAT signaling pathway and uncontrolled hematopoietic cell proliferation^1–4^. Although initially monoallelic, the mutation frequently becomes homozygous through clonal evolution, as mitotic recombination converts the locus via acquired uniparental disomy at 9p and the resulting homozygous clone expands under its selective advantage in the PV marrow^5,6^. While the mutation’s role in promoting erythroid expansion is well established, the specific differentiation stage at which its transcriptional consequences first become biologically manifest has remained poorly characterized.

Normal erythropoiesis proceeds through a well-defined series of morphologically and molecularly distinct stages: megakaryocyte-erythroid progenitors (MEPs) commit to the erythroid lineage and give rise to proerythroblasts, the first morphologically recognizable erythroid-committed cells, which then differentiate into basophilic, polychromatic, and orthochromatic erythroblasts and finally reticulocytes^7,8^. The proerythroblast-to-basophilic erythroblast transition is one of the most molecularly dense steps in erythropoiesis: the cell commits fully to the hemoglobin synthesis machinery, ribosome biogenesis expands dramatically, and survival becomes increasingly dependent on EPO/EPOR→JAK2→STAT5 signaling. The increasing dependence on EPO/EPOR signaling at this transition may explain why the functional consequences of JAK2^V617F^ are particularly pronounced at this stage, since the mutation causes constitutive, EPO-independent JAK2 activation and STAT5 phosphorylation regardless of EPO levels^1,3^, conferring a clonal advantage precisely where EPO dependence becomes the dominant survival signal. The critical role of STAT5 here is underscored by mouse models in which Stat5 is essential for JAK2^V617F^ driven disease^9,10^.

The mutation is present in hematopoietic stem cells and inherited by all downstream progeny, yet the clinical phenotype manifests predominantly as excessive erythrocyte production, raising a fundamental question: at what point along the erythroid trajectory does a genomically present mutation become a transcriptionally active disease driver? This study addressed this question through a multi-layer transcriptomics integration approach combining scRNA-seq DEG analysis, NicheNet ligand-receptor modeling, pseudotime trajectory inference, and inferCNV copy number analysis on the scRNA-seq dataset of Grasshoff and Kalmer^11^ and Kalmer et al.^12^, comprising erythroid colonies grown in colony forming unit assays from peripheral blood mononuclear cells of PV patients and healthy controls, cultured with or without interferon alpha, and further supported by bulk transcriptome data.

## Results

### Data Quality and Cell Type Annotation

Cell type annotations and the erythroid differentiation hierarchy used throughout this study were previously established by Kalmer et al.^12^, encompassing erythroid populations alongside immune populations. Quality control confirmed that these annotations are supported by the transcriptional structure of the data: the elbow plot plateaued after the fifth principal component, per-cell-type distributions of nFeature_RNA, nCount_RNA, and mitochondrial percentage matched expected erythroid biology including the decline of mitochondrial content with maturation, and UMAP or PCA plots stratified by sample, stage, treatment, and condition showed no systematic batch effect, with the first two components resolving biologically meaningful cell type distinctions (Supplementary Fig. S1; Figure 2).

**Figure 1.**
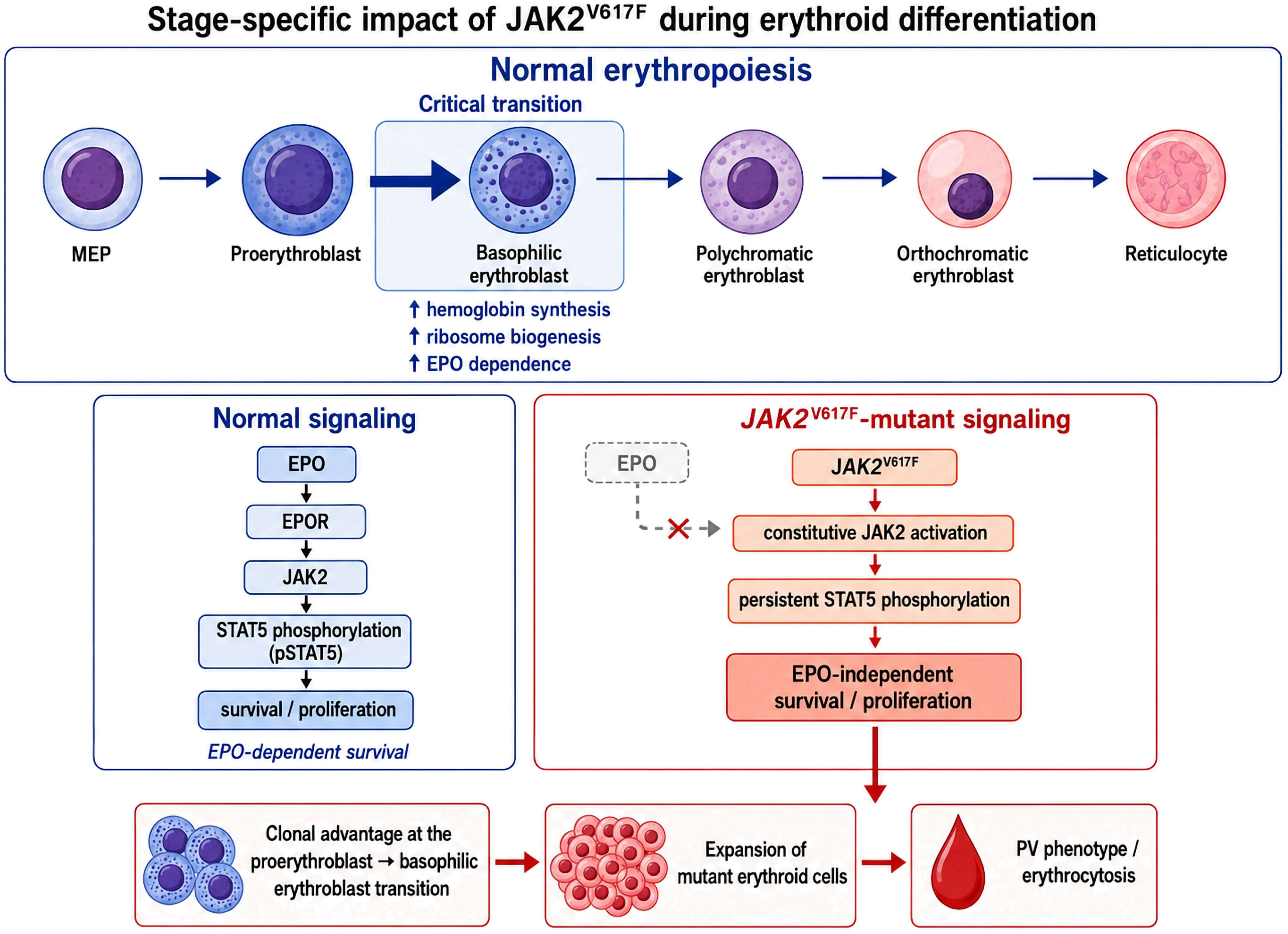
Stage specific impact of JAK2^V617F^ during erythroid differentiation. Top: the normal erythroid cascade from megakaryocyte erythroid progenitors through proerythroblasts, basophilic, polychromatic and orthochromatic erythroblasts to reticulocytes, with the proerythroblast to basophilic erythroblast step marked as the critical transition at which hemoglobin synthesis, ribosome biogenesis and EPO dependence all increase. Bottom left: normal EPO dependent signaling, in which EPO binding to EPOR activates JAK2 and STAT5 phosphorylation and thereby supports survival and proliferation. Bottom right: JAK2^V617F^ mutant signaling, in which JAK2 is constitutively active and STAT5 is persistently phosphorylated independently of EPO. Bottom: the resulting clonal advantage at the proerythroblast to basophilic erythroblast transition, the expansion of mutant erythroid cells, and the erythrocytosis that defines the PV phenotype.

**Figure 2.**
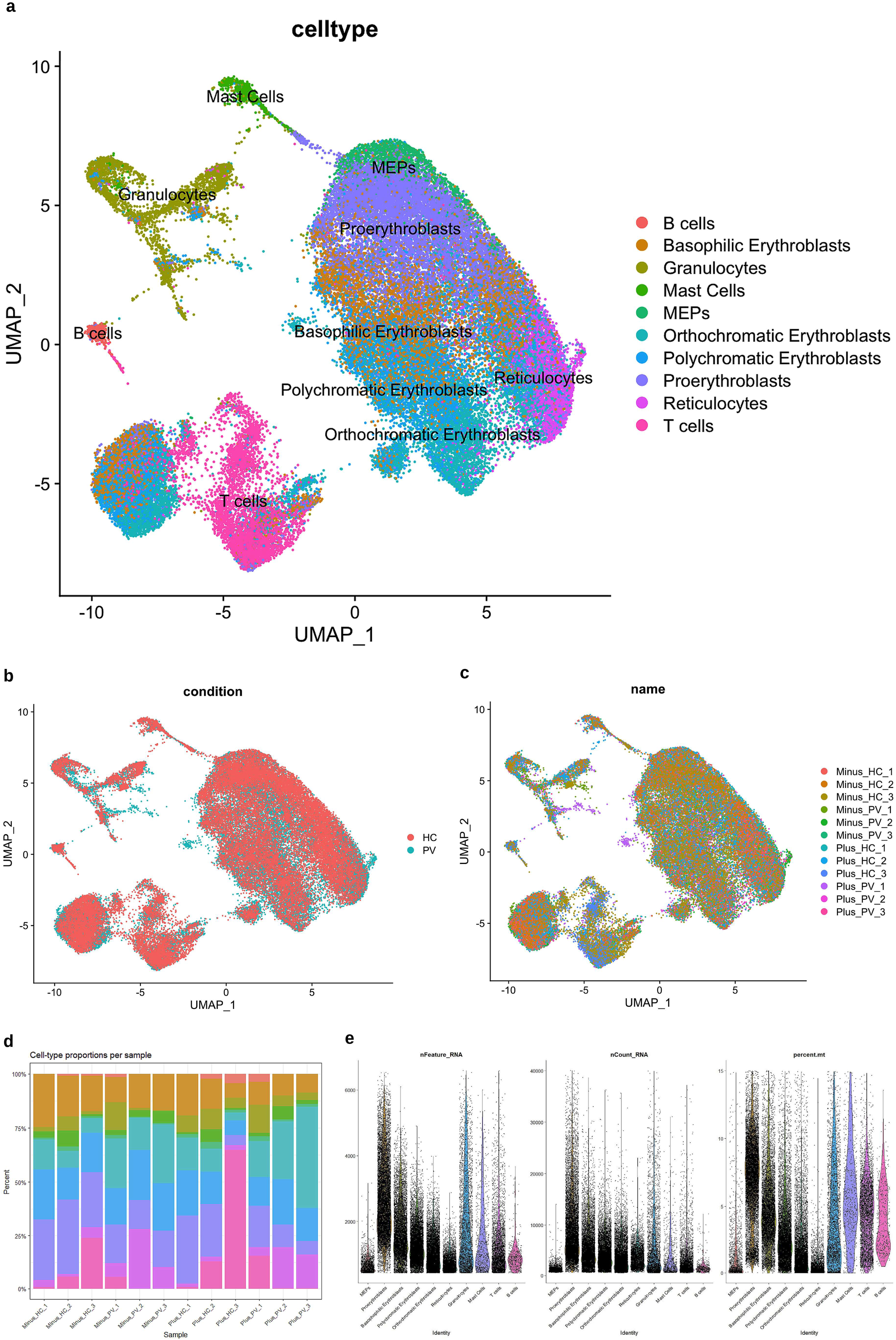
Single-cell transcriptomic overview of healthy control and Polycythemia Vera samples. (a)UMAP of the integrated single-cell RNA-sequencing dataset, with cells colored according to their annotated cell type. Identified populations included B cells, T cells, granulocytes, mast cells, megakaryocyte–erythroid progenitors (MEPs), proerythroblasts, basophilic erythroblasts, polychromatic erythroblasts, orthochromatic erythroblasts, and reticulocytes. (b) UMAP representation of the same cells colored by clinical condition: healthy control (HC) or polycythemia vera (PV). (c) UMAP colored by sample of origin, illustrating the contribution and distribution of individual treated and untreated (Minus and Plus) samples across the integrated dataset. (d) Relative proportions of the annotated cell types within each sample. Each stacked bar represents one sample and is normalized to 100%, color code in (2a) applies (e) Violin plots showing the distributions of the number of detected genes per cell (nFeature_RNA), total RNA counts per cell (nCount_RNA), and percentage of mitochondrial transcripts (percent.mt) across the annotated cell populations.

### Bulk Transcriptome Analysis of PV

Bulk transcriptome profiling of PV versus healthy control samples provided a general landscape of the transcriptional deregulation in PV samples^13^. In below sections, it will be shown that this landscape is an independent line of evidence that converges on the same biological themes identified in the scRNA-seq DEG analysis. Upregulated genes in bulk PV samples with respect to HC samples were enriched for erythrocyte differentiation, gas and oxygen transport, hemoglobin metabolism, and reactive oxygen species metabolic process, directly reflecting the expanded erythroid compartment of PV and consistent with the excess reactive oxygen species produced by JAK2^V617F^ mutant cells that modify the marrow microenvironment and promote genomic instability^14,15^. Downregulated genes were enriched for lymphocyte differentiation, RNA splicing, and B cell activation, reflecting skewing of hematopoiesis toward the erythroid and myeloid lineages at the expense of lymphoid output, in line with the reported impairment of lymphoid differentiation by JAK2^V617F16^. KEGG analysis showed widespread upregulation across the complement and coagulation cascade, consistent with the elevated thrombotic risk that defines the clinical phenotype of PV^17^. Together, these bulk-level signatures reinforce the below cell-type-resolved findings, indicating that independent analytical approaches arrive at a consistent landscape of PV pathophysiology.

### Progressive Transcriptional Disruption Along the Erythroid Lineage

In scRNA data, a stringent DEG intersection strategy separated two complementary signals for each cell type: a disease effect, defined by genes differentially expressed across all PV versus HC comparisons, and a treatment effect, defined by genes consistent across all treated (Plus) versus untreated (Minus) comparisons. Requiring consistent overlap between DEGs across all relevant comparisons including the DEGs from the bulk samples ensured that only the most reproducibly differential expressed genes were carried forward.

Applying a stringent DEG intersection strategy across erythroid cell types revealed progressive transcriptional disruption that intensifies with maturation. Disease affected genes rose from only 7 in MEPs to 56 in proerythroblasts and 53 in basophilic erythroblasts, then 51 in polychromatic erythroblasts, a peak of 103 in orthochromatic erythroblasts, and 45 in reticulocytes. The treatment response showed its own progression, rising from 22 treatment affected genes in MEPs to 39 in proerythroblasts, peaking at 57 in basophilic erythroblasts, then declining through polychromatic (53), orthochromatic (41), and reticulocyte (18) stages. At every stage the treatment response was dominated by a canonical interferon stimulated gene signature including *STAT1*, *IFIT3*, *IFI44L*, *RSAD2*, *IFI27*, *PLSCR1*, and *EPSTI1*. Because the disease effect gene sets were additionally required to intersect the bulk DEGs, cell types contributing few cells, in particular MEPs, are expected to be underrepresented in this ranking.

The disease effect at the MEP stage was minimal, consistent with a population that harbors the mutation but has not yet triggered its transcriptional consequences. By the proerythroblast stage the disease burden was already substantial, with 56 genes: *SNCA* marked the aberrant differentiation trajectory, *MAP2K2* implicated the RAS/MAPK cascade downstream of JAK2^V617F^, *SMARCE1* and the histone variants pointed to chromatin remodeling disruption, and *CDT1* suggested excessive DNA replication licensing consistent with the hyperproliferative phenotype. Yet despite this burden, proerythroblasts did not exhibit the trajectory-level or communicational divergence that defines the disease phenotype, as described in below sections.

The basophilic erythroblast stage is where the JAK2^V617F^ mutation’s transcriptional consequences become most fully manifest, with the highest treatment-affected gene count of any cell type and the highest combined disease and treatment burden among the erythroid progenitor stages. Among its 53 disease-affected genes, the signature is qualitatively distinct from earlier stages. *ERFE* upregulation begins suppressing hepcidin, initiating the iron dysregulation that is a hallmark of PV^18^; *MAP2K2* reflects RAS/MAPK co-activation downstream of mutant JAK2; and *SNCA* marks the aberrant differentiation trajectory of the expanding clone. Epigenetic reprogramming through *JMJD1C*, *KDM7A*, and the histone variants *HIST1H2AE* and *HIST1H4H* begins reshaping chromatin, while *NPM1*, *EIF3A*, *PRPF4B*, and *RBM27* reflect altered ribosome biogenesis, RNA splicing, and translational control, collectively the architecture of a cell type that has transitioned from harboring a silent mutation to actively executing a disease-specific transcriptional program.

The full list of 53 disease-affected genes at this stage comprises: *ABCC4, ACKR1, ADA, ADSL, AGGF1, ANKRD36, AQP3, ARL4A, ATP2B1, BCAT1, BEX1, C2orf88, CD59, CD99, CNIH4, CNOT4, CYSTM1, EIF3A, ERFE, FBXO9, GYPB, GYPE, HACD2, HIST1H1E, HIST1H2AE, HIST1H4H, ITGA4, JAZF1, JMJD1C, KDM7A, MAP2K2, MT2A, NPM1, PAXBP1, PIP5K1B, PLEK2, PQLC1, PRPF4B, PTP4A1, RAB8B, RAPGEF6, RBM27, RNPC3, RPL3, RSRP1, SEC62, SLC14A1, SMARCE1, SNCA, SRGN, STK38, STOM*, and *USP12*.

### Cellular Communication Evidence via NicheNet Analysis

At the earliest erythroid stage, NicheNet revealed no condition-dependent restructuring of intercellular communication: across all six pairwise comparisons the ligand-receptor interaction potentials in MEPs were essentially unchanged, indicating that neither PV nor interferon alpha alters the prioritized repertoire of this bipotent progenitor (Figure 5a,b). Differential ligand expression was nonetheless detectable, with S100A8 upregulated across erythroblast and lymphoid populations in untreated samples (Minus_PV versus Minus_HC), LGALS9 and HLA-A upregulated under both treatment comparisons irrespective of disease status, and the angiogenesis inhibitor COL18A1 downregulated in disease. The absence of any qualitative shift in ligand-receptor interaction potential despite these ligand-level fluctuations marks the MEP as a population that carries the mutation without yet engaging a disease-specific communication program.

**Figure 3.**
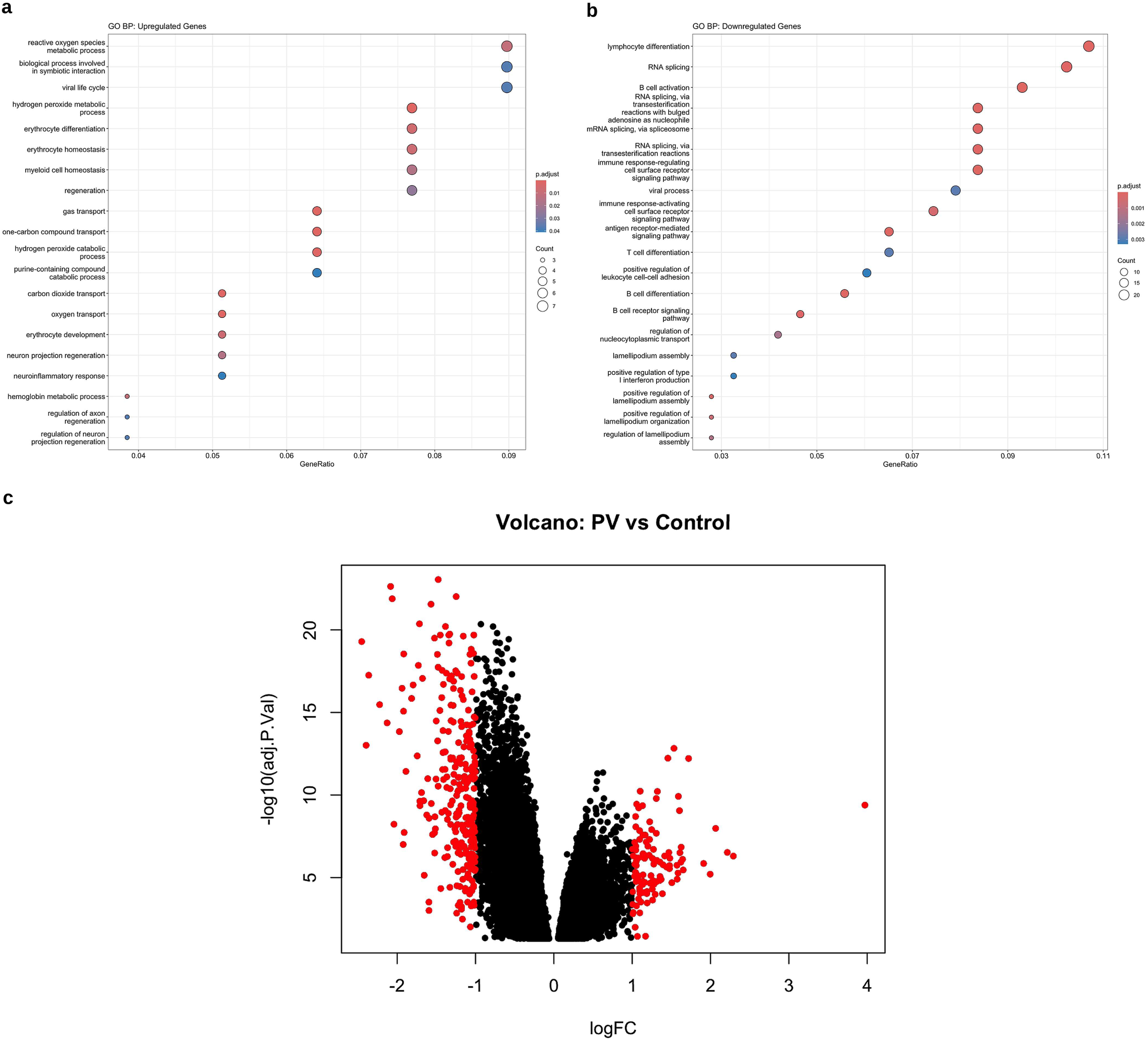
Bulk transcriptome comparison of PV versus healthy control. (a) GO term enrichment for the upregulated genes, dominated by reactive oxygen species metabolism, erythrocyte differentiation, gas and oxygen transport, and hemoglobin metabolism. (b) GO term enrichment for the downregulated genes, enriched for lymphocyte differentiation, RNA splicing, and B cell activation. (c) Volcano plot; 324 differentially expressed genes were identified, with 89 upregulated and 235 downregulated in PV.

**Figure 4.**
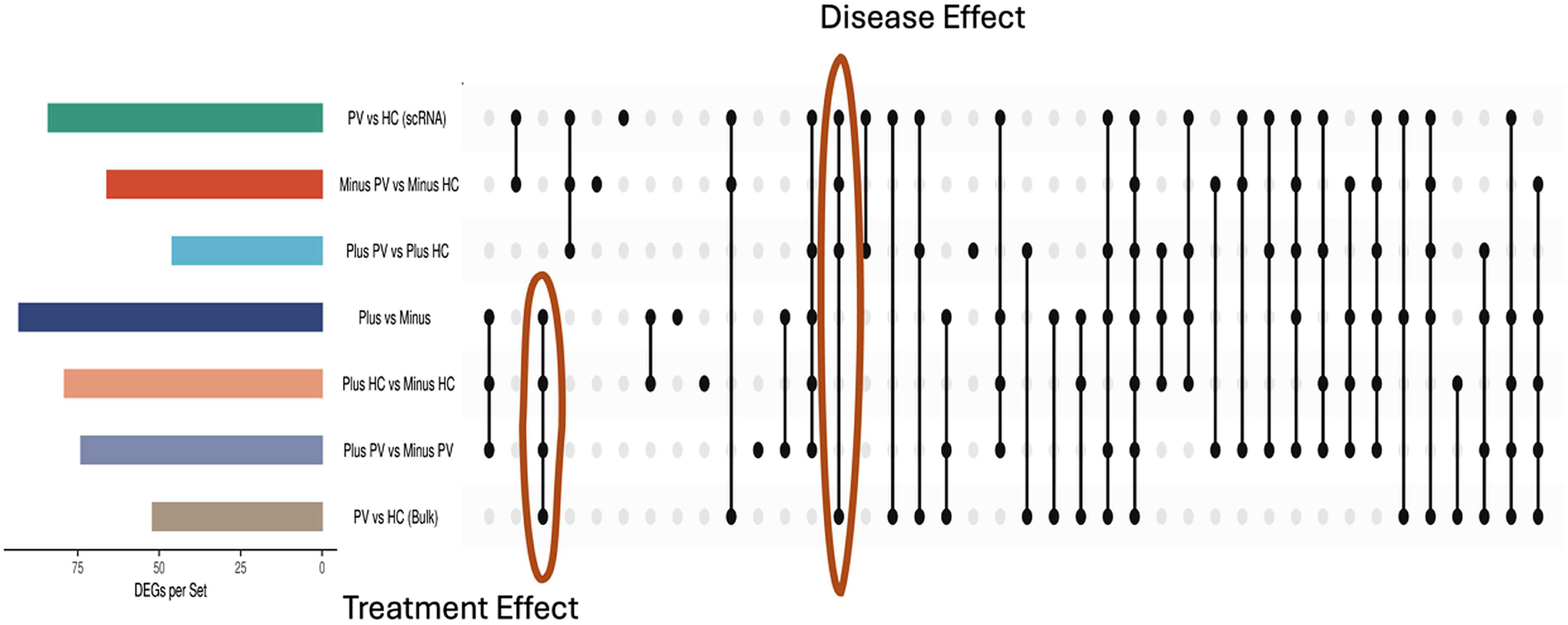
UpSet plot illustrating the DEG intersections used to define the treatment effect and the disease effect gene sets for each cell type. The complete gene lists for all cell types are given in Supplementary Table S2.

**Figure 5.**
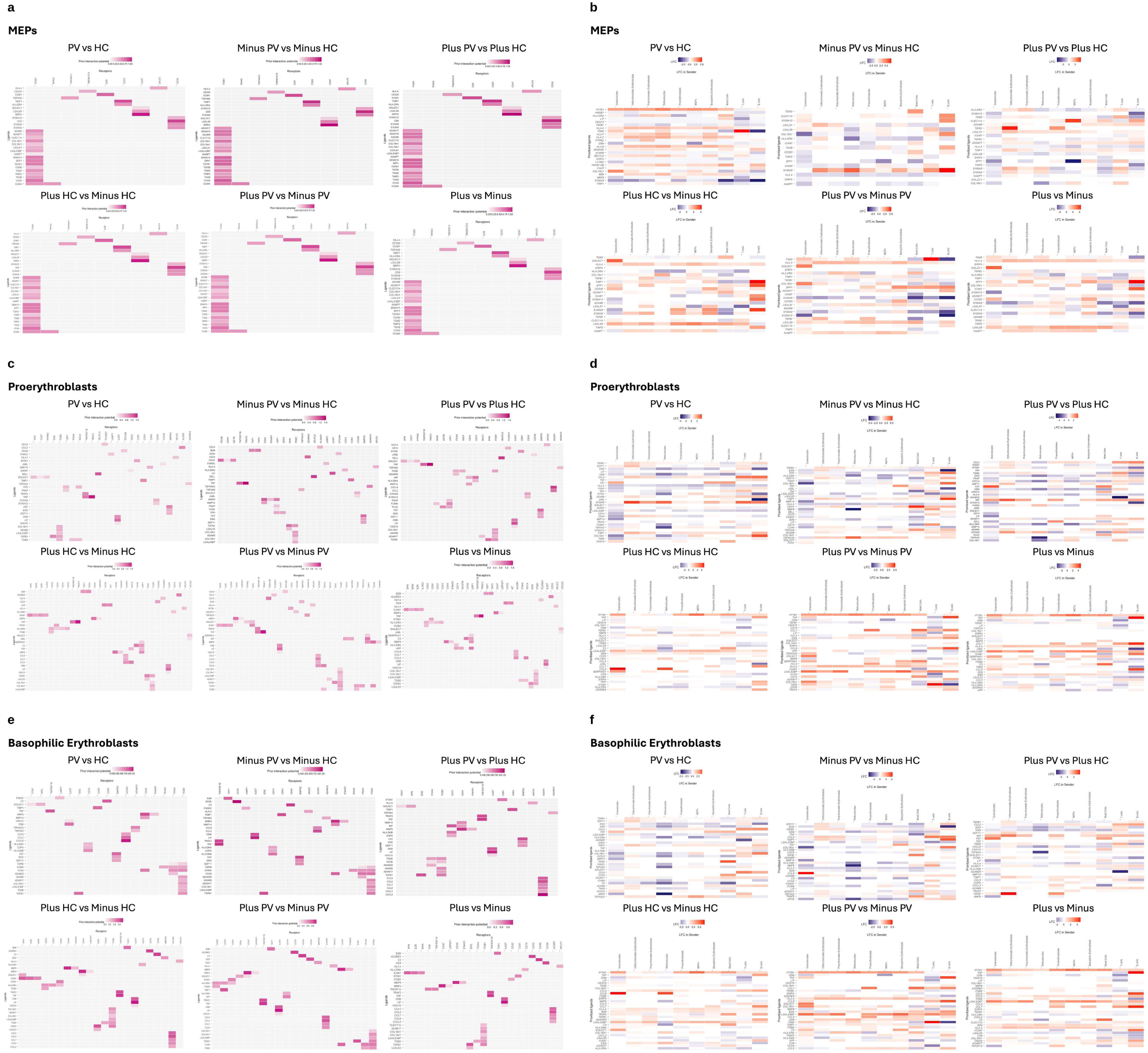
NicheNet ligand-receptor interaction analysis for MEPs, proerythroblasts, and basophilic erythroblasts across six pairwise comparisons. (a) Interaction potentials and (b) ligand expression log fold change in sender populations for MEPs. (c) Interaction potentials and (d) ligand expression log fold change in sender populations for proerythroblasts. (e) Interaction potentials and (f) ligand expression log fold change in sender populations for basophilic erythroblasts.

By the proerythroblast stage a coherent disease-associated communication signature begins to take shape, although it remains confined to the TGFβ superfamily and inflammatory cytokine axes of the baseline erythroid repertoire (Figure 5c,d). TNF-TNFRSF1B emerged among the strongest prioritized pairs in the disease comparisons, consistent with elevated TNF signaling in the PV marrow^19^, while ITGB1 supported progenitor niche retention through integrin-mediated adhesion to stromal VCAM1. Additional disease-effect ligands included FAM3C, CD44, MMP14, SIGLEC1, FST, ENG, GDF11, and TGFB1, with SELL appearing specifically at this stage to suggest selectin-mediated homing, and FCN1 uniquely engaging CD34/PTPRC in untreated disease to indicate early complement lectin pathway activation. The TGFβ superfamily ligands TGFB1, GDF11, ENG, and FST were broadly upregulated in disease. The treatment response engaged a distinct set, including TRAF2-mediated NF-κB/JNK activation, coagulation-associated SERPINC1, inhibitory SIGLEC7, and chemokines CCL2/8 and CXCL3/8 seen in Plus_PV versus Plus_HC but not in the overall comparison, indicating that the chemokine layer is unmasked only under treatment. These signals represent the establishment of the baseline TGFβ axis rather than the chemokine and matrix-remodeling expansion that defines the basophilic erythroblast stage.

NicheNet indicated that basophilic erythroblasts express a qualitatively distinct communication repertoire in PV relative to both earlier and later stages (Figure 5e,f). The TGFβ superfamily axis established earlier (TGFB1-ACVR1/ENG, FST, ENG, GDF11) solidifies here and is now joined by an expanded repertoire including FAM3C, SIGLEC1, TNF, MMP9, MMP14, VCAM, ADAM28, ADAM17, CCL2, CCL5, CCL7, and CCL8. MMP9 and MMP14 upregulation indicates an acquired matrix remodeling capacity, consistent with the aberrant progenitor expansion described in PV^20^, while the TNF-TNFRSF1B axis reinforces the inflammatory environment that sustains clonal growth^19^. The expansion of CCL chemokines as both disease and treatment response ligands specifically at this stage indicates that basophilic erythroblasts acquire a chemokine expression capacity in PV. Because these cells were profiled as colonies in culture rather than in situ, this represents a cell intrinsic potential for immune recruitment rather than a demonstrated remodeling of an intact marrow niche.

In the treatment comparisons, ICAM1-ITGB1/ITGB2 interactions emerged among the strongest pairs, reflecting enhanced adhesive crosstalk in the stimulated disease state consistent with JAK2^V617F^-driven integrin upregulation, and connecting the DEG-level observation of ITGA4 dysregulation to a functional adhesion phenotype detected by an independent method. By contrast, the MEP stage showed no qualitative shift, and the proerythroblast stage engaged only the baseline TGFβ axis without the chemokine and matrix-remodeling expansion described above, further supporting that the cells at MEP-to-proerythroblast transition do not have the disease phenotype. The later erythroid stages sustain these axes rather than replace them: polychromatic erythroblasts add BMP9, LGALS1, LGALS3BP, VCAN, ADAM28, TGFBI and TGM2 to the disease repertoire, engaging CD47, SPN, ITGB1, TAP1 and IFNAR1 (Supplementary Figure S2); orthochromatic erythroblasts reinforce the SIRPA, SIGLEC1 and S100A axes with broad upregulation of CCL2, ADAM17, TGM2 and VCAN (Supplementary Figure S3); and reticulocytes engage CD99-PILRA and PSEN1 interactions specific to terminal maturation while TGFBI, ADAM9 and LGALS3BP remain upregulated (Supplementary Figure S4). The repertoire expansion first seen at the basophilic erythroblast stage is therefore carried forward through terminal maturation rather than re-established at any later stage.

### Pseudotime Trajectory Evidence: The Basophilic Erythroblast Is the Turning Point

Pseudotime analysis, performed independently for each erythroid cell type and colored by stage (Minus_HC, Minus_PV, Plus_HC, Plus_PV), revealed distinct transcriptional continuum profiles for each cell type, in agreement with previous results. In MEPs and proerythroblasts, the four conditions are indistinguishable along the trajectory; however at the basophilic erythroblast stage they cluster into six distinct, biologically coherent regions. From this stage onward, the transcriptional separation is sustained but its underlying mechanism shifts, as the cell-cycle-coupled disease program transitions into apoptotic and stress-dominated programs, identifying the basophilic erythroblast as the precise point at which JAK2^V617F^ becomes a trajectory-altering disease driver.

All condition based pseudotime panels capture the same canonical erythroid maturation ordering (Figure 6a-d). In all four conditions the reticulocytes occupy a terminal end of the principal graph, indicating that the ordering reflects genuine maturation rather than batch– or condition-specific structure. The conditions differed not in the sequence of stages but in the continuity of the graph: in two of the four conditions the cells formed a single connected erythroid manifold in which the MEP-to-proerythroblast step was supported by a contiguous ridge of cells, whereas only the PV conditions are fragmented into multiple partitions with the MEP cluster fully detached in Minus_PV. Non-erythroid populations formed independent components throughout, confirming that the graph did not over-connect unrelated lineages.

**Figure 6.**
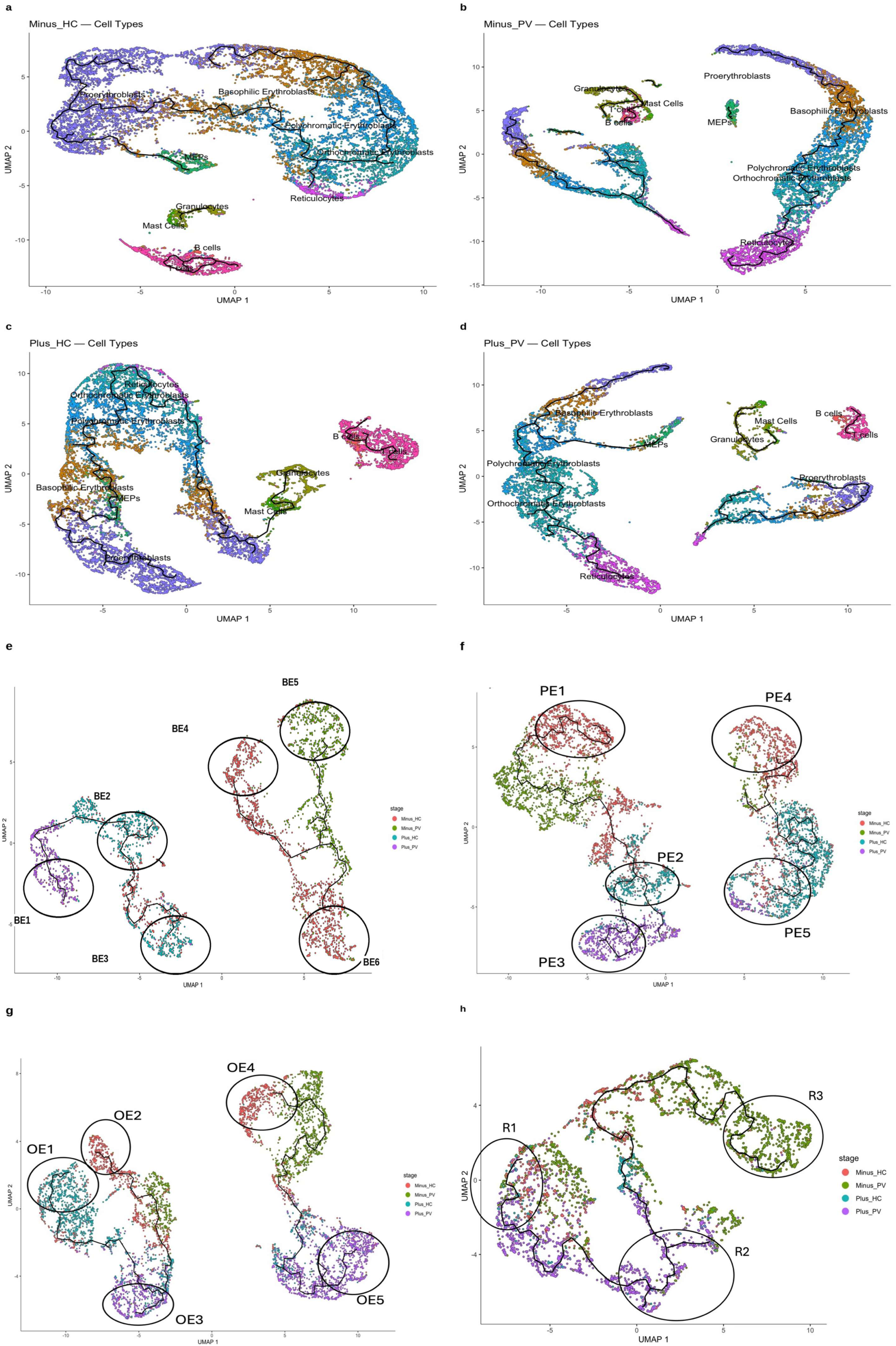
Erythroid differentiation trajectories across the erythroid lineage. (a-d) UMAP embeddings of single-cell transcriptomes colored by annotated cell type, with the Monocle3 trajectory curve overlaid in black: (a) Minus_HC, (b) Minus_PV, (c) Plus_HC, (d) Plus_PV. Each panel was embedded and graph-learned independently; axis scales and pseudotime are therefore internally normalized and not comparable across panels. (e-h) Cell type based pseudotime trajectories, colored by stage (Minus_HC, Minus_PV, Plus_HC, Plus_PV) with condition-enriched subregions circled. (e) Basophilic erythroblasts, where the trajectory first separates into six condition-enriched subregions (BE1 to BE6). (f) Polychromatic erythroblasts (PE1 to PE5), where the Plus_PV subregion loses cell cycle enrichment. (g) Orthochromatic erythroblasts (OE1 to OE5). (h) Reticulocytes (R1 to R3).

Cell type based pseudotime results (Supplementary Figs. S5 and S6, Figure 6e-h; region level enrichment in Supplementary Figs. S7 to S25) indicated that prior to the basophilic erythroblast in the maturation cascade, there is no trajectory effect despite the mutation. MEPs showed no condition-dependent structure along their pseudotime trajectory, cells from all four conditions are distributed uniformly (Supplementary Figure S5). Proerythroblasts, despite already carrying a substantial 56 gene disease signature in the DEG analysis, likewise showed no trajectory wise differences (Supplementary Figure S6). The contrast is the central observation of this section: gene level perturbation alone is insufficient to produce a disease phenotype gradient along a trajectory.

Unlike the two preceding erythroid stages, which showed no condition specific structure, the basophilic erythroblast is the earliest stage at which the conditions diverge along the trajectory itself (Figure 6e). Interpreted by condition, the six regions reproduce the pathway signatures expected from PV biology and from interferon alpha exposure. The untreated PV clone (Minus_PV) establishes the disease baseline in BE5, which is enriched for translation, mitochondrial oxidative phosphorylation, the ER-phagosome pathway, and mitochondrial protein degradation, while interferon signaling is present but weak, ranking below these pathways. Mitochondrial translation is itself a determinant of terminal erythroid maturation and iron homeostasis^21^, so this metabolic profile is what would be expected of an expanding clone that has not been exposed to interferon. The interferon exposed conditions superimpose a strong interferon response on this background. In interferon exposed healthy controls (Plus_HC), BE2 and BE3 display interferon signaling without cell cycle enrichment: BE2 is restricted to interferon alpha signaling, rRNA processing, and amino acid metabolism, whereas BE3 additionally recruits interferon gamma signaling, lymphocyte activation, and antigen presentation. In interferon exposed PV (Plus_PV), BE1 co-enriches interferon signaling with cell cycle and DNA replication programs, combining the JAK2^V617F^ driven proliferative signature with the interferon response. The contrast is informative: the interferon alpha response is engaged in both health and disease, but only in PV is it superimposed on an active proliferative program.

Following the transition point, the disease associated molecular signature persists, while the underlying transcriptional program undergoes a marked shift (Figure 6f). In the interferon exposed healthy controls, PE2 and PE5 (Plus_HC) retain strong interferon alpha, beta, and gamma signaling. The Plus_PV enriched PE3 region no longer shows cell cycle enrichment, instead displaying broad activity in interferon signaling, NF-κB activation, amino acid biosynthesis, apoptotic signaling, and the cytosolic DNA sensing pathway. Among the untreated healthy controls, PE1 (Minus_HC) notably includes negative regulation of type I interferon signaling, which suggests active suppression of the response, and PE4 (Minus_HC) shifts almost entirely to metabolic terms (rRNA processing, TCA cycle, propanoate and lipoic acid metabolism, amino acid metabolism). No region at this stage was enriched for Minus_PV. This shift from a cell cycle coupled program at the basophilic stage to a cell cycle decoupled, apoptosis and stress dominated program marks the exit of these cells from the proliferative phase and, with it, the point beyond which they are no longer susceptible to the proliferation dependent action of interferon alpha.

At the orthochromatic stage the trajectory diversifies into five thematically distinct regions reflecting the diverging requirements of terminal maturation (Figure 6g). The Plus_HC enriched OE1 region diverges toward membrane lipid metabolism, lipid biosynthesis, ganglioside metabolism, Fas signaling, and post-Golgi vesicle transport, consistent with the membrane remodeling required during enucleation, in which lipid raft organization, Rac GTPase dependent cytoskeletal polarization, and vesicle trafficking drive nuclear extrusion and reticulocyte formation^22,23^; the concurrent Fas signal may reflect a negative autoregulatory axis constraining erythroid expansion under stress^24,25^. The Plus_PV enriched OE3 region shows weaker enrichment dominated by Huntington disease pathway, cell cycle, chromosome maintenance, EIF2AK1/HRI response to heme deficiency^26^, and negative regulation of ferroptosis^27^, while the Plus_PV enriched OE5 region is defined by proteolysis, NRF2 antioxidant detoxification, FOXO mediated transcription^28^, autophagy, mitophagy, and TORC2 signaling. The Minus_HC enriched OE2 and OE4 regions maintain strong interferon signaling with NF-κB, MHC class I antigen processing, and RIG-I signaling^29^, and uniquely in OE4, positive regulation of erythrocyte differentiation and RORA/B/C circadian regulation^30^; interferon induces antigen processing programs through NLRC5 and CIITA^31–33^, which is consistent with this pattern. As at the polychromatic stage, no region was enriched for Minus_PV. The disease imprint here is no longer proliferative but metabolic, oxidative, and stress adaptive.

Reticulocytes complete the lineage with a single trajectory running from a heterogeneous Plus_PV enriched R1 region through a Plus_PV oriented R2 region to a Minus_PV enriched R3 region (Figure 6h), all dominated by translational and mitochondrial themes that reflect the post-nuclear transcriptome focused on hemoglobin synthesis. Taking the conditions in turn, the Minus_PV enriched R3 region is the most strongly enriched of the three and shows interferon alpha signaling, asparagine N-linked glycosylation, PINK1-PRKN mitophagy, ferroptosis regulation, lysosomal acidification, and IL12 induced JAK-STAT signaling, a profile aligned with iron dependent oxidative stress and compensatory mitochondrial clearance^34–36^. The Plus_PV oriented R2 region shows translation as its top term alongside the ER-phagosome pathway, 43S complex formation, mitochondrial protein degradation, NADH dehydrogenase complex assembly, erythrocyte homeostasis, and macroautophagy, and R1 adds porphyrin metabolism, oxidative phosphorylation, JAK-STAT signaling, and TP53 regulation of metabolic genes. No region at this stage was enriched for either healthy control condition. The oxidative and mitophagy signature of the untreated PV reticulocytes suggests that terminal maturation in PV proceeds under sustained iron dependent oxidative stress^37^.

The cumulative pseudotime evidence converges on a single finding: across the entire erythroid lineage there is a single sharp boundary, at the basophilic erythroblast stage. Before this stage, JAK2^V617F^ is genomically present but trajectory silent; at it, the mutation engages the cell cycle and resolves the trajectory into six condition-enriched regions for the first time; after it, the imprint is sustained across polychromatic, orthochromatic, and reticulocyte trajectories but shifts from proliferation toward stress, apoptosis, and metabolic adaptation as cells exit cycle. The basophilic erythroblast is therefore not merely the first cell type to register the disease, but the precise point at which the diseased niche becomes mechanistically distinct from the normal erythroid program.

### InferCNV Evidence

InferCNV analysis allowed us to identify and score the regions of increased and decreased expression intensity. The analysis revealed a consistent increase along the erythroid lineage in the percentage of cells whose mean absolute CNV score exceeds the reference derived threshold of 0.0257 (Figure 7a; Supplementary Fig. S26), from 3.16% in MEPs to 6.21% in proerythroblasts, 6.15% in basophilic erythroblasts, 7.18% in polychromatic erythroblasts, 13.87% in orthochromatic erythroblasts, and a peak of 21.03% in reticulocytes. This progressive increase along the erythropoiesis cascade suggests that the effect of the JAK2 mutation becomes clearer as cells differentiate into more mature states.

**Figure 7.**
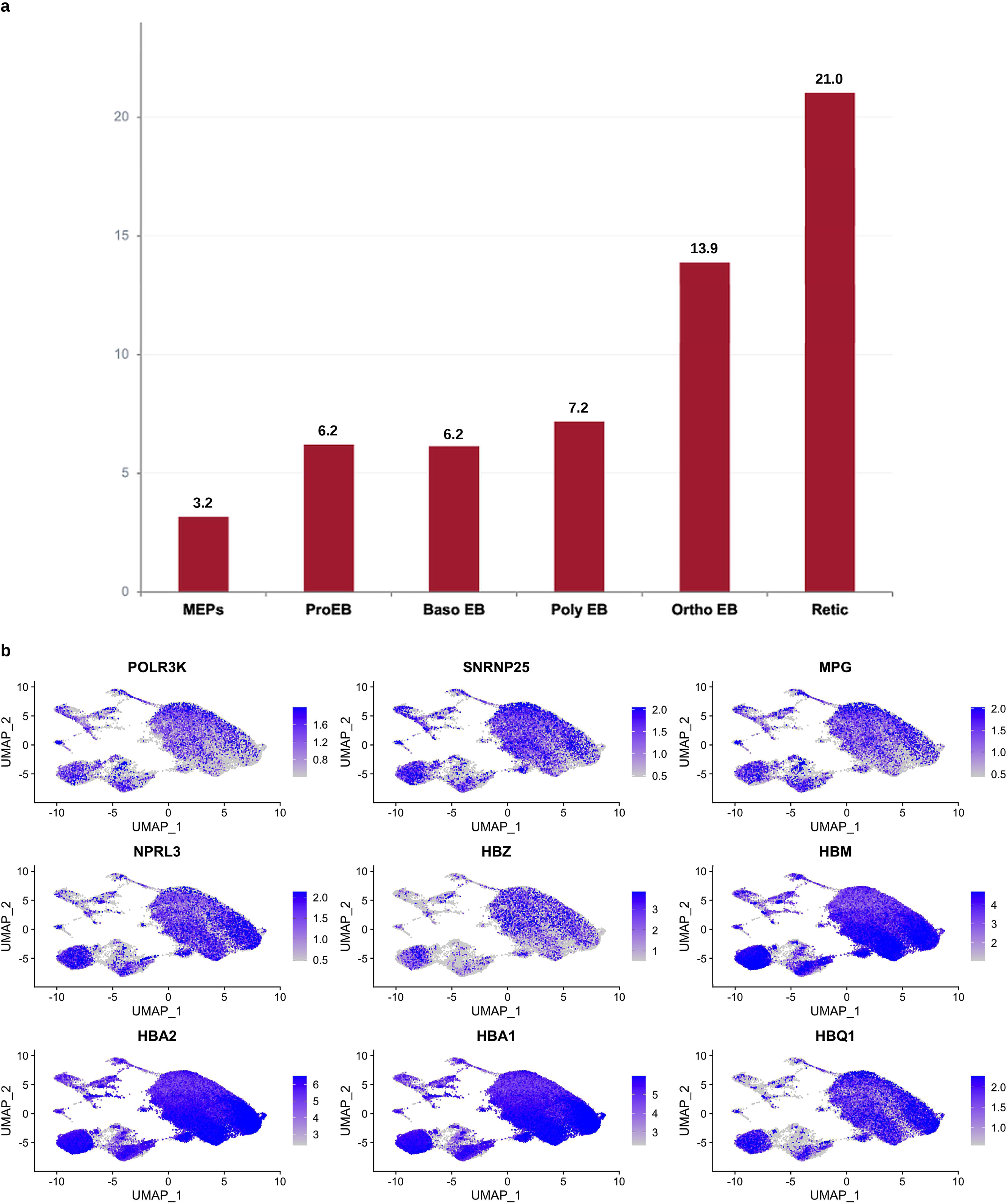
InferCNV copy number variant analysis. (a) CNV burden percentages for each inferCNV run across the erythroid differentiation order, showing a progressive increase from MEPs to reticulocytes. (b) Feature plots for the genes most strongly correlated with the CNV score.

The genes most strongly correlated with the CNV score, *POLR3K*, *SNRNP25*, *MPG*, *NPRL3*, *HBZ*, *HBM*, *HBA2*, *HBA1*, and *HBQ1*, all cluster at chromosome 16p13.3, the alpha-globin gene cluster and its flanking region. *HBZ*, *HBM*, *HBA2*, *HBA1*, and *HBQ1* are the alpha-globin genes themselves and because *NPRL3*, *MPG*, *POLR3K*, and *SNRNP25* are immediate genomic neighbors, their shared alteration likely reflects a single regional CNV event rather than nine independent gene-specific signals.

Although the CNV burden percentage at the basophilic erythroblast stage (6.15%) is comparable to that of proerythroblasts (6.21%), the CNV data are informative rather than contradictory when read alongside the other modalities. The near equivalence of CNV burden percentages between the two stages, against a complete absence of pseudotime trajectory divergence in proerythroblasts, indicates that copy number alterations detectable by inferCNV precede the transcriptional trajectory divergence visible by pseudotime. This temporal layering is coherent: the inferred copy-number alterations act as upstream events that progressively reshape the transcriptome, and it is at the basophilic erythroblast stage that this accumulated transcriptional instability crosses a threshold sufficient to produce a detectable communication profile.

### The Proerythroblast to Basophilic Erythroblast Transition

Examining the genes altered genes at the basophilic erythroblast stage relative to proerythroblasts revealed an unexpected decline in *KIT*, *STAT5A*, *CDK6*, *CASP3*, and *SMARCC1* (Figure 8). Feature plots show that, except for *CASP3* and *SSMARCC1*, these genes are expressed predominantly in earlier erythroid populations and lose expression as cells mature, therefore their coordinated downregulation at the transition marks the molecular switch at which the disease signature first consolidates. The simultaneous loss of a chromatin remodeling scaffold (*SMARCC1*), the direct JAK2^V617F^ effector (*STAT5A*), a cell cycle driver (*CDK6*), and the apoptotic executioner (*CASP3*) at the same transition motivates the *SMARCC1*/BAF155 upstream regulator hypothesis described below.

**Figure 8.**
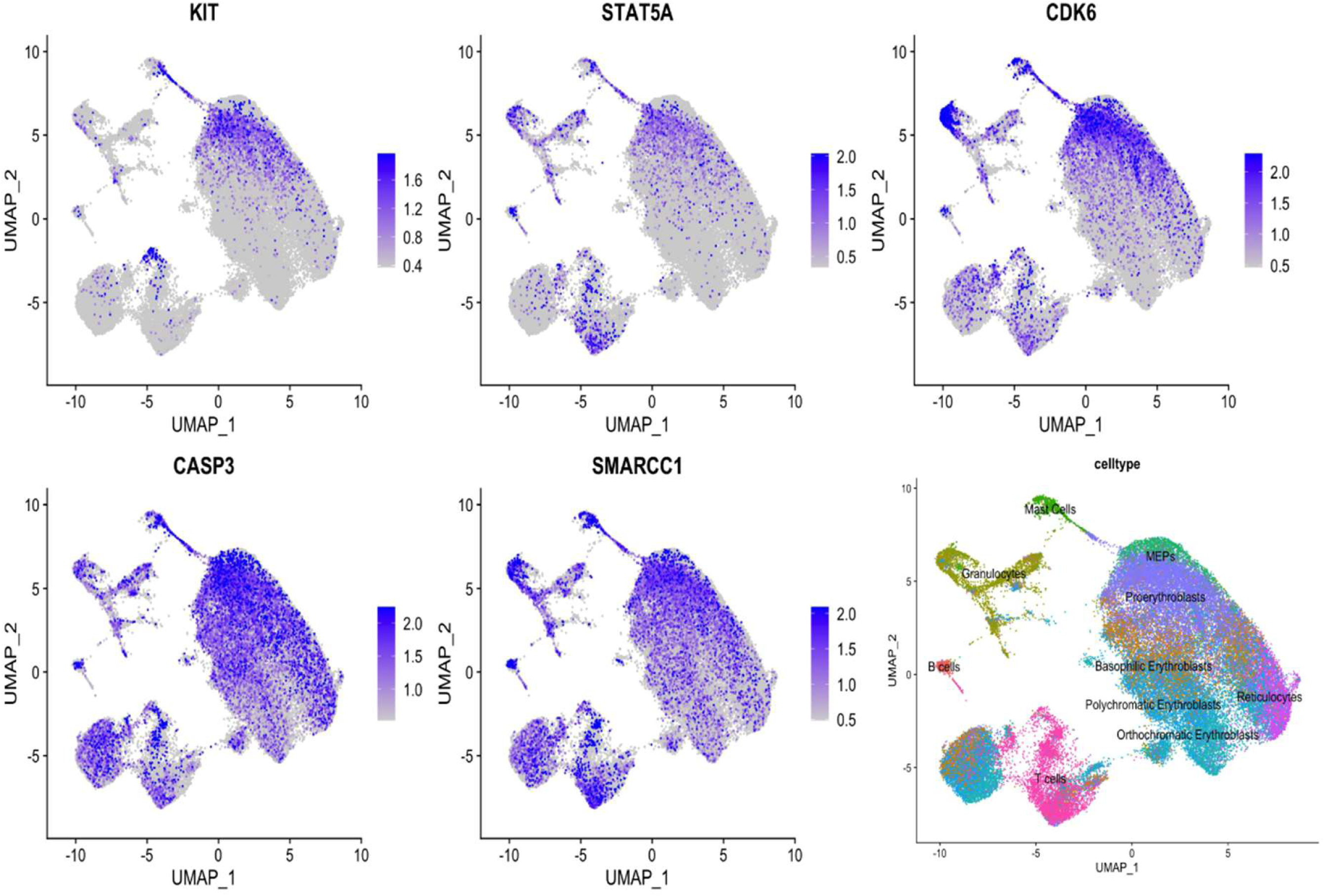
Feature plots for the downregulated genes in basophilic erythroblasts with respect to proerythroblasts, shown alongside the cell type UMAP for reference.

When the treatment effect is considered, interferon alpha shows its broadest intervention at this stage, simultaneously triggering the ISG cascade (*STAT1*, *IFIT3*, *IFI44L*, *SAMD9L*), suppressing translation through *EIF2AK2* and multiple initiation factors, and inducing pro-apoptotic *TNFSF10* (TRAIL), identifying basophilic erythroblasts as being at the cellular state in which disease associated clonal expansion is most pronounced and treatment induces the strongest opposing transcriptional response^12^. Disease activation and treatment counterresponse reaching their combined maximum in the same population establishes the basophilic erythroblast as the principal therapeutic battleground of PV. Contrasting this stage with the immediately preceding proerythroblast stage isolates the molecular events specifically activated upon entry into the basophilic state, pinpointing the transcriptional switch that marks the onset of PV pathology.

The DEG analysis at the proerythroblast-to-basophilic erythroblast transition identified *SMARCC1* (BAF155), a core structural subunit of the SWI/SNF chromatin remodeling complex, as downregulated together with *CASP3*, *CDK6*, and *STAT5A*, and therefore as a potential upstream master regulator of their coordinated loss. Importantly, this occurs while *GYPA*, *GYPB*, *SLC4A1*, *HBA1*, *HBA2*, and *HBB* are concurrently upregulated across the same transition, indicating that the withdrawal of proliferative and survival signaling accompanies, rather than opposes, the onset of globin synthesis and membrane-skeleton assembly. *SMARCC1* is an essential scaffolding subunit of the E-RC1 (EKLF Coactivator-Remodeling Complex 1), which cooperates with GATA-1 and EKLF/KLF1 to open chromatin at critical erythroid genes such as the β-globin locus^38^. Erythroid cells are especially sensitive to disruption of this system because *SMARCA2* (BRM), the paralog that compensates for the loss of *SMARCA4* (BRG1) in SWI/SNF complexes in other lineages. Accordingly, *SMARCA4* hypomorphic mutation arrests erythroid differentiation at the basophilic to polychromatic erythroblast transition^39^, and conditional BAF155 knockout produces cytopenia with failure of hematopoietic regeneration through impaired chromatin priming^40^.

In the context of JAK2^V617F^ PV, *SMARCC1* downregulation at this transition could function as an upstream switch: loss of the chromatin remodeling scaffold would reduce accessibility at the regulatory regions of *CASP3*, *CDK6*, and *STAT5A* simultaneously, providing a single cause for a coordinated multi-gene effect. This model remains a hypothesis rather than a demonstrated mechanism, since chromatin accessibility was not assayed here and differential expression alone cannot establish the causal relationship.

### Targets and Candidates for Drug Repurposing

Following the basophilic erythroblast stage, the polychromatic pseudotime regions showed the emergence of NF-κB signaling, apoptotic activation, and cytosolic DNA sensing, indicating that replicative stress initiated at the basophilic stage has escalated to DNA damage sensing responses. The orthochromatic stage, with its peak of 103 disease affected DEGs and the emergence of nuclear extrusion machinery disruption, represents the terminal amplification of transcriptional disruption building since the basophilic erythroblast stage.

The sharp decline in interferon alpha responsive genes, from 57 in basophilic erythroblasts to 18 in reticulocytes, reflects the progressive loss of transcriptional responsiveness to treatment as cells approach terminal differentiation, consistent with the clinical observation that interferon alpha reduces but rarely fully eliminates the malignant clone. Interferon alpha debulks the clone at its roots, triggering apoptosis in early progenitors from MEPs to basophilic erythroblasts while leaving terminal maturation intact, hence normal erythropoiesis continues as malignant cell numbers fall; it induces JAK/STAT activation and preferentially eliminates ribosome-high PV subclones^12^. The progressive loss of interferon alpha responsiveness in orthochromatic erythroblasts and reticulocytes, combined with continued and even amplified disease-associated disruption at these stages, identifies late erythroid differentiation as the principal escape window of the PV clone from interferon alpha mediated cytoreduction.

The convergence of disease-associated molecular alterations at the basophilic erythroblast stage was centered on MAP2K2-driven RAS/MAPK signaling, ERFE-mediated iron dysregulation, NPM1 dysregulation, epigenetic reprogramming through JMJD1C and KDM7A, and the coordinated downregulation of CASP3, CDK6, and STAT5A. The findings identify several rational drug repurposing opportunities from the existing approved and clinical-stage pharmacopeia.

The established therapeutic backbone of PV provides the foundation for any combination strategy targeting this stage. The coordinated loss of CASP3, CDK6 and STAT5A at this transition dismantles the survival and apoptotic checkpoints of the maturing erythroblast, and normal expression of these genes would favor apoptosis over the expanded erythroid output that characterizes PV. The therapeutic aim at this stage is therefore to restore that apoptotic competence. These transcripts also decline as part of normal maturation, so the disease specific component is their behavior relative to the JAK2 wild type cells of the same patients^12^, and the agents below are proposed based on the programs they engage rather than on transcript abundance alone. Ruxolitinib, a JAK1/2 inhibitor acting downstream of JAK2^V617F^, achieves hematocrit control and symptom benefit in PV and myelofibrosis^41^. Ropeginterferon alfa-2b is approved for PV and acts through activating the interferon program, which in the present analysis elicited its most extensive opposing transcriptional response at the basophilic erythroblast stage^42^. In newly diagnosed PV, the combination of ruxolitinib with interferon alpha was shown to achieve 92 percent peripheral blood count remission at 24 months, 60 percent molecular remission and a reduction in median JAK2^V617F^ allele burden from 47 to 7 percent, and it induced CASP3 while also acting on STAT5 and CDK6, thereby engaging three of the four convergence point targets identified here^43^.

Building on this backbone, the stage-specific vulnerabilities suggest additional targets. MAP2K2 upregulation reflects RAS/MAPK co-activation downstream of JAK2^V617F^, a parallel proliferative signal not addressed by JAK inhibition alone, whose importance is supported by the dependence of JAK2^V617F^ PV progenitors on DUSP1 to withstand inflammatory and DNA-damage stress while sustaining proliferation^44^. ERFE upregulation, first appearing here, marks the earliest actionable iron-regulatory target in PV^18^. STAT5A is the direct downstream effector of JAK2^V617F^, strictly required for JAK2^V617F^ driven polycythemia in mouse models^9,10^. CASP3 dysregulation reflects suppression of the apoptotic executioner, which in healthy erythropoiesis is required at precisely this maturation window for enucleation^22,23^; Navitoclax, a BH3 mimetic that neutralizes BCL-2, BCL-XL and BCL-W and thereby lowers the threshold for executioner caspase activation, is mechanistically matched to this defect: it restores intrinsic apoptosis downstream of the suppressed executioner rather than requiring CASP3 transcript to be restored. It has been evaluated in solid tumors and, in combination with ruxolitinib, in a phase 3 myelofibrosis trial against best available therapy (NCT04468984), where thrombocytopenia is the principal dose limiting toxicity^45,46^.

A multi-node intervention model targeting the basophilic erythroblast stage consolidates these opportunities on the existing interferon alpha or ruxolitinib backbone: acting on the primary JAK2^V617F^→STAT5A axis, closing off the RAS/MAPK proliferative escape, restoring iron homeostasis disrupted by ERFE, and reactivating the suppressed apoptotic executioner CASP3 all at the single differentiation stage where these vulnerabilities converge, before the disease accumulates the additional gene-level disruptions seen at the orthochromatic erythroblast stage where treatment efficacy is already largely lost.

## Discussion

This study converges across multiple independent analytical modalities on a unified conclusion: while PV is initiated by the JAK2^V617F^ mutation in early hematopoietic progenitors, its transcriptional consequences do not manifest in a biologically meaningful and cell type-specific manner until the basophilic erythroblast stage, making this population the pivotal node of disease initiation along the erythroid differentiation trajectory.

The mechanistic basis for this stage-specific activation is rooted in the biology of erythroid differentiation. The proerythroblast-to-basophilic erythroblast transition is the point where the cell switches its survival dependency fully to EPO/JAK2; before the transition it can survive through multiple pathways, but after basophilic erythroblast commitment JAK2 becomes the dominant gatekeeper, which is exactly when a constitutively active JAK2 begins to provide a measurable clonal advantage. The strict requirement for STAT5 in JAK2^V617F^ driven polycythemia in mouse models^9,10^ reinforces this hypothesis. Hence, we suggest that the convergence of disease-associated alterations at this stage, centered on MAP2K2-driven RAS/MAPK signaling, ERFE-mediated iron dysregulation, NPM1 dysregulation, and epigenetic reprogramming through JMJD1C and KDM7A, presents the earliest actionable disease signature in PV.

The pseudotime evidence is particularly compelling: the complete absence of trajectory divergence in proerythroblasts, despite their substantial 56 gene disease signature and comparable CNV burden to basophilic erythroblasts, demonstrates that transcriptional perturbation alone is insufficient to produce a trajectory-level disease phenotype. It is at the basophilic erythroblast stage that the accumulated transcriptional, communicational, and genomic alterations collectively cross a threshold that produces a detectably altered differentiation trajectory.

The NicheNet findings add a critical intercellular signaling dimension. The solidification of the TGFβ superfamily and matrix remodeling axis specifically at the basophilic erythroblast stage, accompanied by the expansion of CCL chemokines and MMP9/MMP14 upregulation, indicates that the disease phenotype extends beyond the intracellular transcriptional program to the ligand repertoire these cells are able to present^19,20^. The basophilic erythroblast is therefore not merely a passive recipient of accumulated molecular damage but a potential active participant in shaping its microenvironment. Because the profiled cells were derived from colony cultures, this inference is limited to expression capacity and awaits confirmation in intact marrow.

The therapeutic implications are substantial: interferon alpha induces its broadest opposing transcriptional response at the basophilic erythroblast stage, which also represents the final developmental stage at which treatment efficacy remains maximal. The progressive loss of responsiveness in later populations, combined with the continued amplification of disease-associated disruption, creates a narrowing therapeutic window optimally targeted at the basophilic erythroblast stage. The drug repurposing analysis identifies multiple agents that could be rationally combined on a backbone of interferon alpha or ruxolitinib to simultaneously address the vulnerabilities converging at this stage. Addressing the *SMARCC1*/BAF155 disruption is also promising: If the coordinated downregulation of *CASP3*, *CDK6*, and *STAT5A* is driven by a single upstream chromatin remodeling deficit, strategies that restore SWI/SNF function could correct all three downstream deficiencies at once. However, no agent currently upregulates SMARCC1 transcription, hence restoration of the chromatin remodeling scaffold remains an unmet therapeutic need and a priority for future drug discovery.

Several limitations of the analyses presented here should be noted. The single cell data were generated from erythroid colonies grown in vitro from peripheral blood mononuclear cells, with interferon alpha added to the culture, hence the samples report a cell intrinsic response to interferon alpha rather than the response of an intact marrow to systemic therapy, and the ligand receptor inferences describe expression capacity rather than intercellular signaling demonstrated in situ. Disease effect gene sets were additionally required to intersect the bulk DEGs, so cell types represented by few cells, in particular MEPs, are likely to be underrepresented. The SMARCC1 hypothesis rests on differential expression alone, since chromatin accessibility was not assayed in the samples. Finally, the drug repurposing candidates are computational inferences from transcriptional signatures and require experimental validation.

Nevertheless, these findings provide a mechanistic framework for understanding PV pathogenesis, identify stage-specific marker genes with potential diagnostic utility, and offer a rational basis for the development of targeted combination therapies aimed at the basophilic erythroblast stage which is the precise moment when JAK2^V617F^ transitions from a silent passenger to an active disease driver.

## Methods

### Dataset and Preprocessing

The primary scRNA-seq dataset was retrieved from Grasshoff and Kalmer^11^, deposited at Zenodo, and described in Kalmer et al.^12^. The dataset comprises single cell transcriptomes of erythroid colonies generated in colony forming unit assays from peripheral blood mononuclear cells of polycythemia vera patients and healthy controls, cultured for ten days with 500 U/mL interferon alpha-2b added to the methylcellulose (Plus) or omitted (Minus), yielding four experimental groups: Plus_PV, Minus_PV, Plus_HC, and Minus_HC, enabling six pairwise comparisons that isolate the disease effect, the treatment effect, and their interaction. Cell type annotations established by the original authors were used, encompassing erythroid populations (MEPs, proerythroblasts, basophilic erythroblasts, polychromatic erythroblasts, orthochromatic erythroblasts, and reticulocytes) alongside immune populations (granulocytes, T cells, B cells, and mast cells). Because interferon alpha was applied to the colony cultures rather than administered to patients, the Plus conditions report a cell intrinsic response to interferon alpha rather than the response of an intact marrow to systemic therapy. The dataset was assembled, integrated and annotated by the original authors using Seurat version 4.2.0. All analyses reported here were performed in R using Seurat version 5.0^47^.

### Quality Control and Batch Effect Assessment

Quality control of the dataset, which resulted in 54,307 cells and 19,820 genes, was performed prior to all downstream analyses. UMAP projections were stratified by condition, treatment, stage, and patient to assess batch effects. PCA was performed to evaluate whether transcriptional variance reflected biological cell type distinctions rather than technical artifacts. Elbow plot analysis was used to determine the number of principal components capturing the dominant sources of transcriptional variance. Cells were filtered to retain those with mitochondrial gene percentage below 15 percent, more than 400 detected features and fewer than 40,000 mRNA counts.

### Differential Gene Expression Analysis

Differentially expressed genes were identified across six pairwise comparisons for each cell type. A stringent intersection strategy was applied: genes differentially expressed across all PV versus HC comparisons were retained as disease-affected genes, while genes consistent across all Plus versus Minus comparisons were retained as treatment-affected genes, isolating the transcriptional effect of treatment independently of disease context. Single cell differential expression was performed with the FindMarkers function of Seurat using a Wilcoxon rank sum test, and genes were retained at an average absolute log fold change above 0.5 with a Benjamini-Hochberg adjusted p value below 0.05. For the bulk microarray data, comprising 41 PV and 21 control samples, differential expression was performed with limma (version 3.54.0, R 4.2.2) at an absolute log fold change above 1.0 with an adjusted p value below 0.05. In both strategies, requiring consistent overlap across all relevant comparisons and validating against bulk DEGs reduced false positives and ensured that only the most reproducible signals were carried forward for each cell type. Bulk transcriptome analysis was performed for differential expression, followed by GO and KEGG pathway enrichment, with pathway-level integration and visualization performed using Pathview^48^ and functional enrichment using Metascape^49^.

### NicheNet Ligand-Receptor Interaction Analysis

NicheNet^50^ was used to model intercellular communication by predicting ligand-receptor interactions between sender and receiver cell populations. Six pairwise comparisons were performed for each receiver cell type across the four conditions. The analysis identified upstream ligands from sender populations whose downstream target genes were differentially expressed in the receiver population. Ligand activity scores, ligand-receptor interaction potentials, and ligand expression log fold changes in sender populations were computed to characterize the niche remodeling axis at each erythroid stage.

### Pseudotime Trajectory Analysis

Monocle3^51^ was used to perform pseudotime trajectory inference on each erythroid cell type and condition independently. Each of the four condition-specific datasets was embedded and graph-learned independently rather than on a jointly integrated object; consequently, UMAP coordinates and pseudotime values are internally scaled within a panel and are not directly comparable between panels. Comparisons across conditions were therefore restricted to graph topology, namely the ordering of annotated stages along the principal graph, the continuity or fragmentation of that graph, and the position of branch points. For each cell type, cells were embedded in reduced-dimensional space, a principal graph was fitted, and pseudotime values were assigned.

Trajectory plots were colored by condition (stage: Minus_HC, Minus_PV, Plus_HC, Plus_PV) to assess whether condition-dependent trajectory divergence was present. Where a trajectory was observed within a cell type across conditions, the most differentiated regions were selected and subjected to a region-based DEG and pathway enrichment analysis to characterize the biological processes driving the separation.

### InferCNV Analysis

InferCNV^52^ is used to infer large-scale chromosomal copy number alterations or regions of differential gene expression intensity from gene expression data. B cells, T cells, granulocytes and mast cells, drawn from both interferon exposed and unexposed samples, served as the reference population for the cell type specific runs; no reference cells were supplied for the whole object run, for which the package uses the mean of all cells as the baseline. A mean absolute CNV score threshold was defined as the maximum of the reference cell median plus three median absolute deviations and the reference cell 99th percentile, giving 0.02570775, and the percentage of cells exceeding this threshold was computed for each erythroid cell type. Genes most strongly correlated with the CNV score were identified by Spearman correlation between gene expression and the per cell CNV score across all cells, retaining genes with a mean absolute correlation coefficient above 0.13, and their chromosomal locations were mapped to detect recurrent genomic alterations associated with the JAK2^V617F^ clone.

### Drug Repurposing Analysis

Disease-affected genes identified at the basophilic erythroblast stage were mapped to known drug targets using the existing approved and clinical-stage pharmacopeia. Candidate drugs were evaluated based on their mechanism of action relative to the identified molecular, current clinical development status, and existing evidence in myeloproliferative neoplasms. A multi-node intervention model was constructed to identify rational combination strategies targeting the converging vulnerabilities at this single differentiation stage.

## Ethics declarations

This study is a secondary computational analysis of previously published publicly available, de-identified human transcriptomic datasets (Zenodo 10.5281/zenodo.14204623; GEO GSE26049). No new human samples were collected. Ethical approval and informed consent for the original sample collection were obtained by the investigators of the source studies, as reported in Kalmer et al. No additional ethical approval was required for the present analysis.

## Data availability

The single-cell RNA sequencing dataset analyzed in this study is publicly available at Zenodo (https://doi.org/10.5281/zenodo.14204623). The bulk transcriptome dataset is available from the Gene Expression Omnibus under accession number GSE26049. The disease effect and treatment effect gene sets for every cell type are provided in Supplementary Table S2.

## Author contributions

D.K. designed and performed the computational analyses and wrote the manuscript. P.P. designed the analyses, conceived and supervised the study and revised the manuscript. Both authors reviewed and approved the final manuscript.

---

**Supplementary Figure S7.**
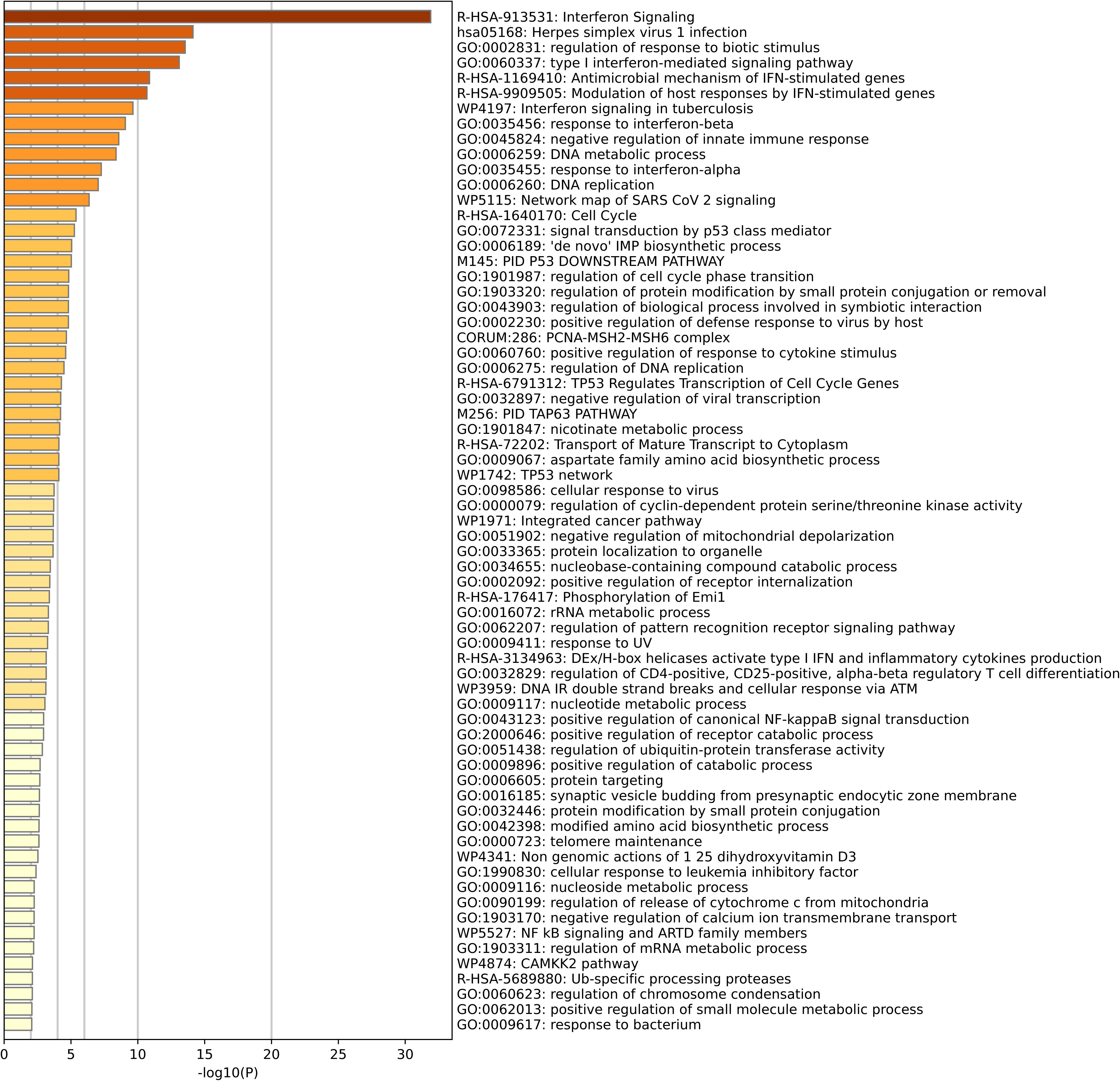
Pathway enrichment for region BE1, basophilic erythroblasts.

**Supplementary Figure S8.**
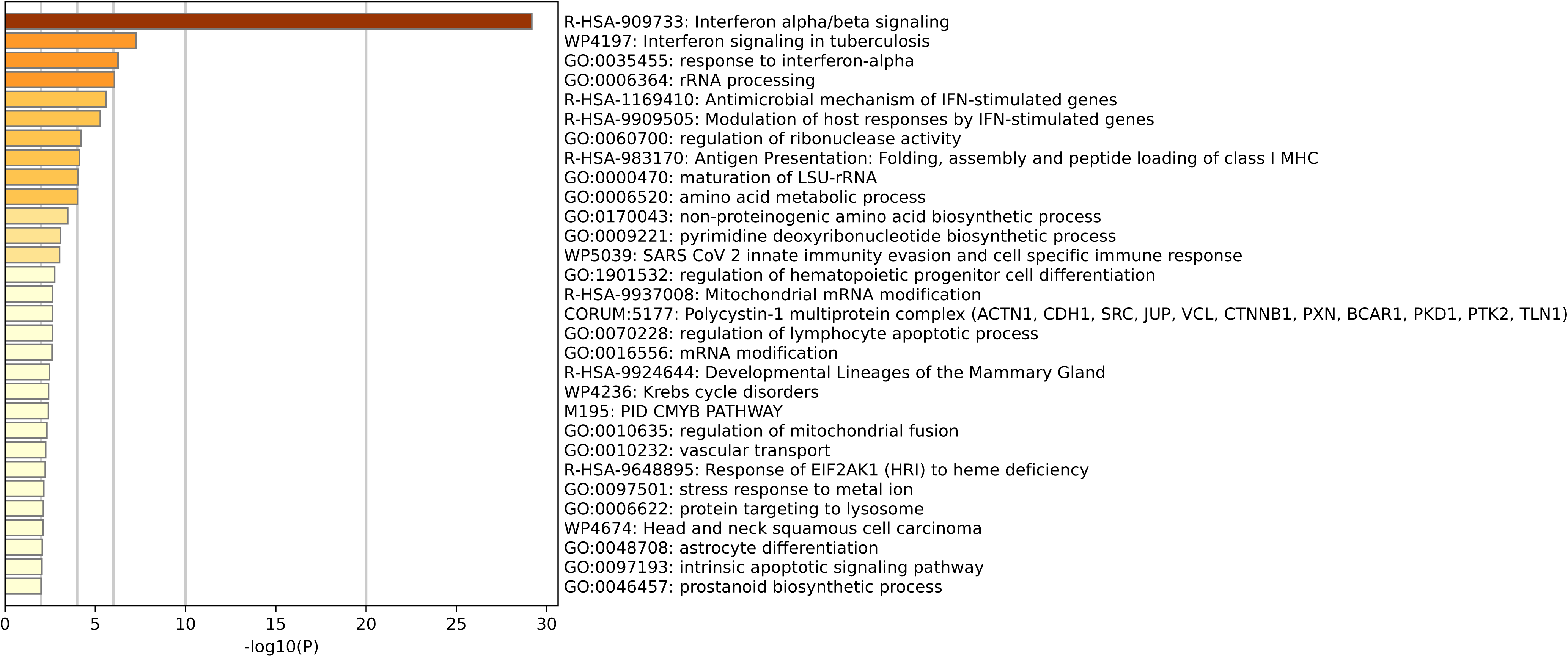
Pathway enrichment for region BE2, basophilic erythroblasts.

**Supplementary Figure S9.**
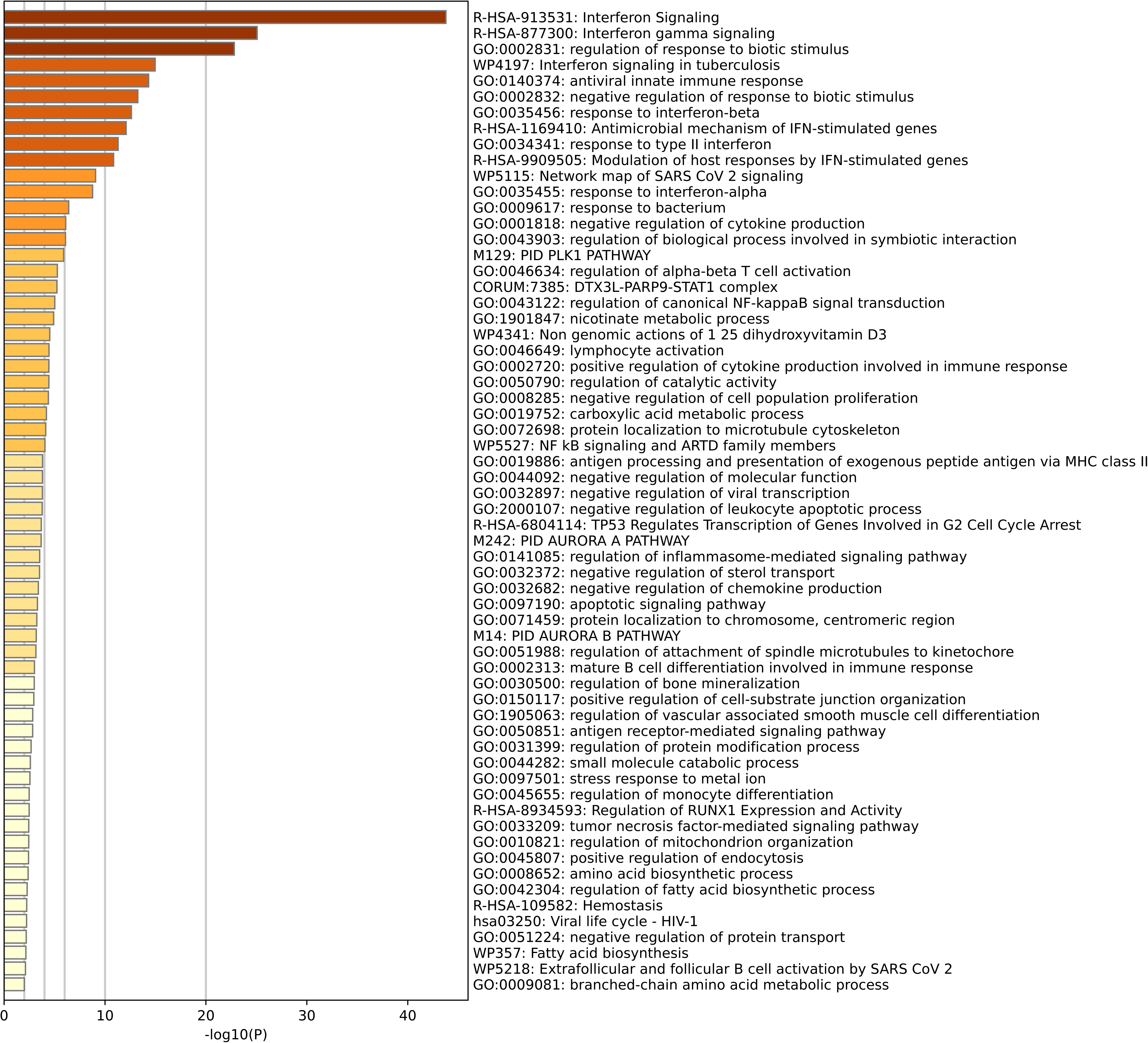
Pathway enrichment for region BE3, basophilic erythroblasts.

**Supplementary Figure S10.**
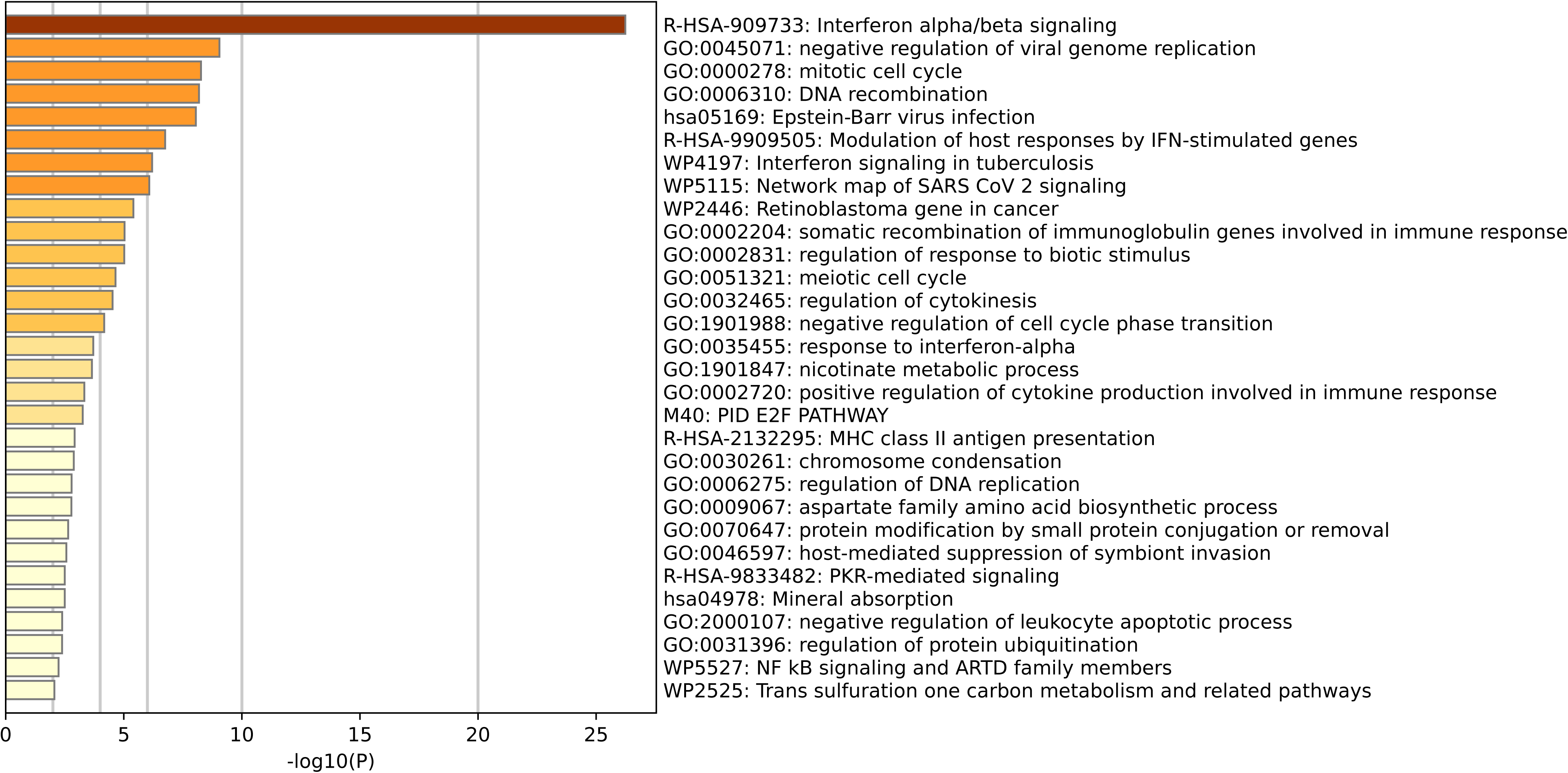
Pathway enrichment for region BE4, basophilic erythroblasts.

**Supplementary Figure S11.**
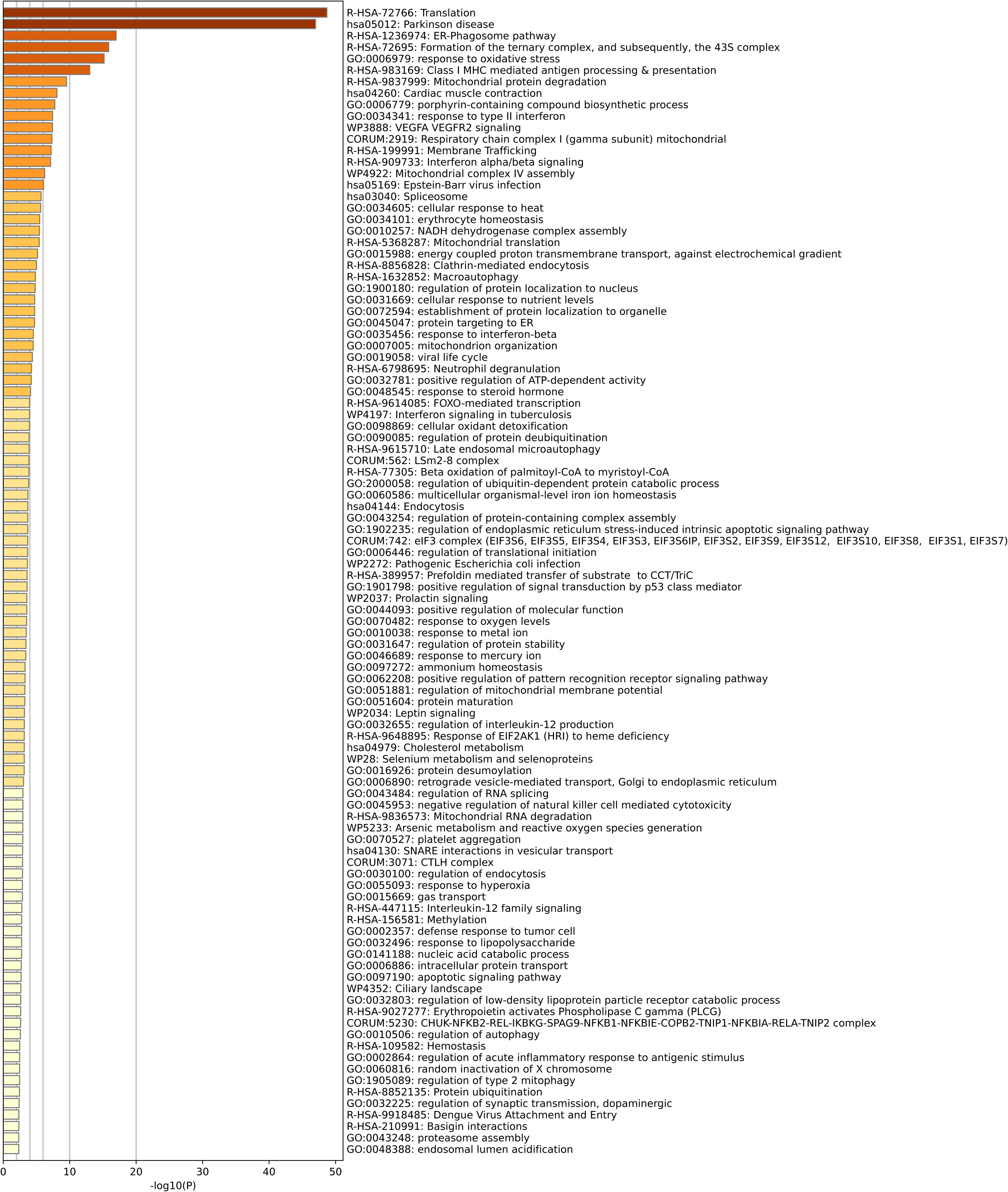
Pathway enrichment for region BE5, basophilic erythroblasts.

**Supplementary Figure S12.**
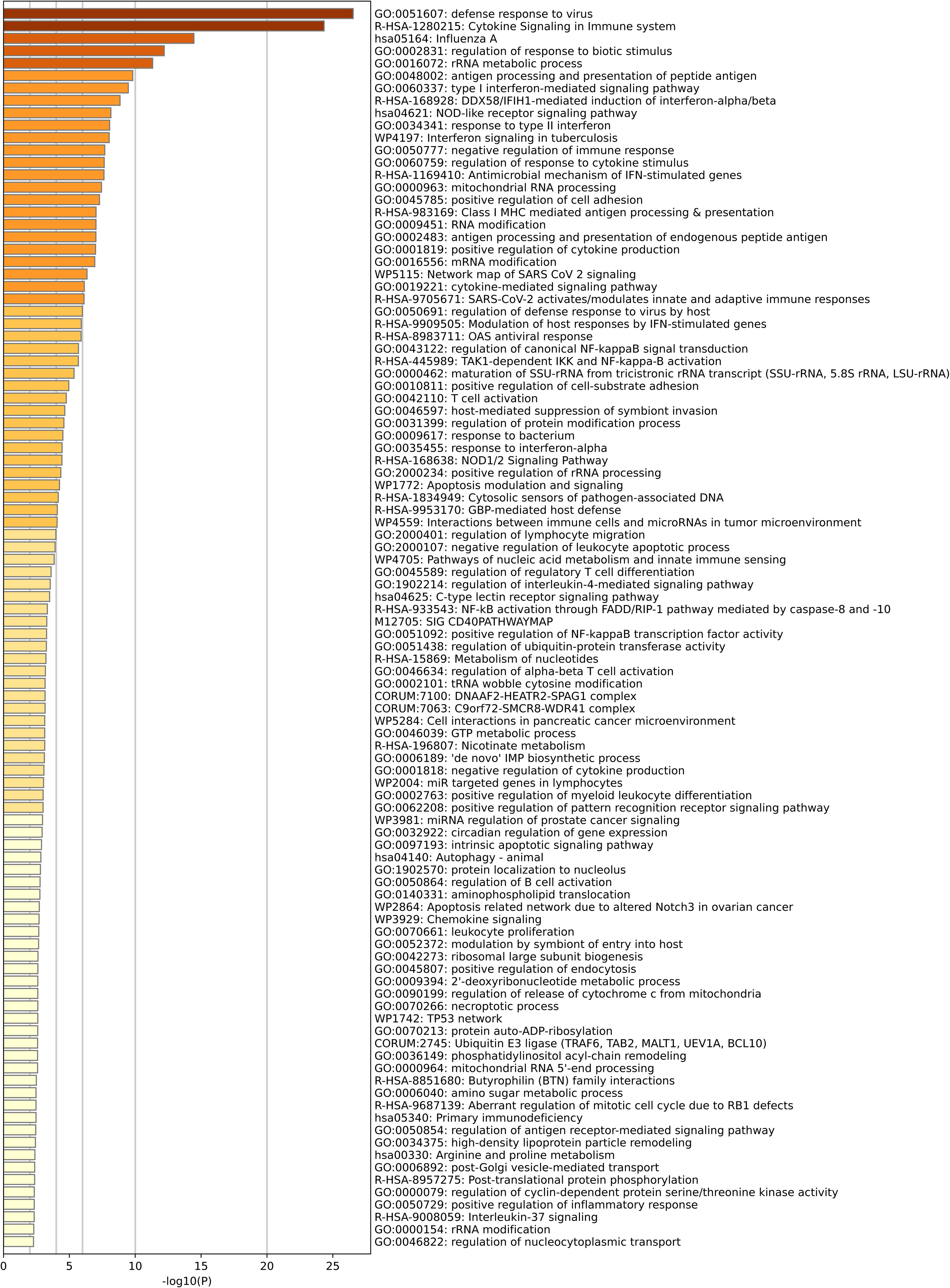
Pathway enrichment for region BE6, basophilic erythroblasts.

**Supplementary Figure S13.**
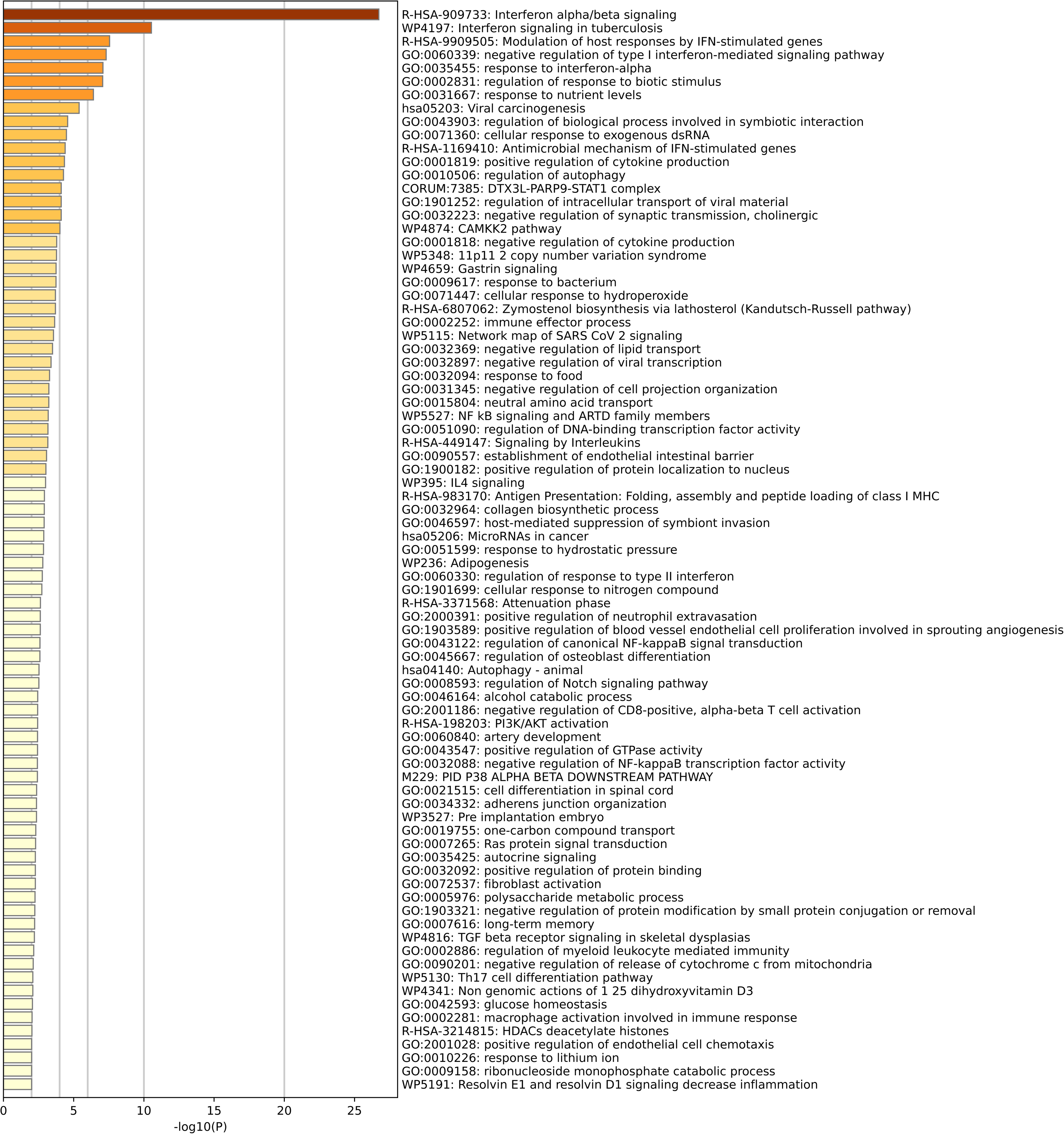
Pathway enrichment for region PE1, polychromatic erythroblasts.

**Supplementary Figure S14.**
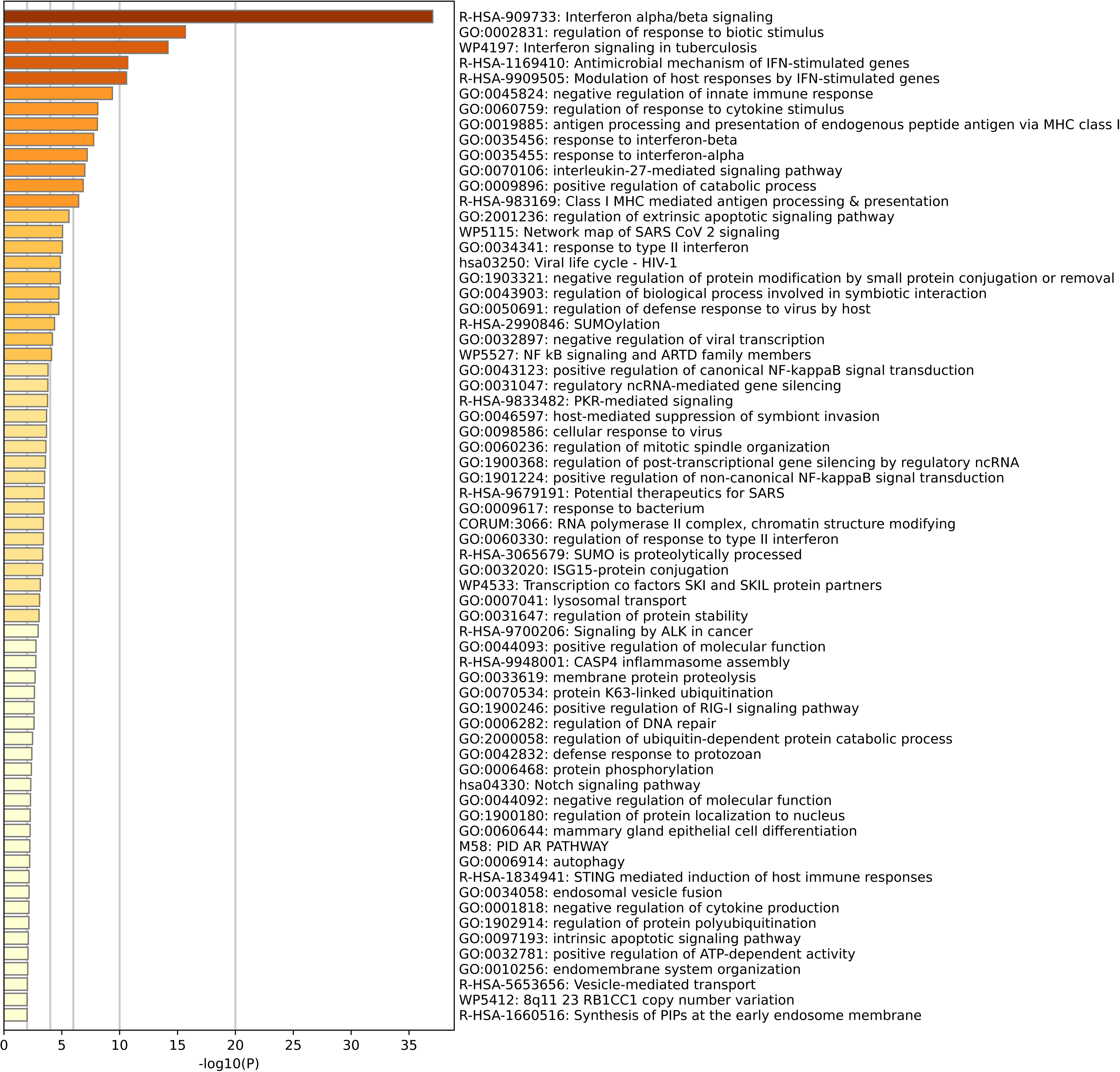
Pathway enrichment for region PE2, polychromatic erythroblasts.

**Supplementary Figure S15.**
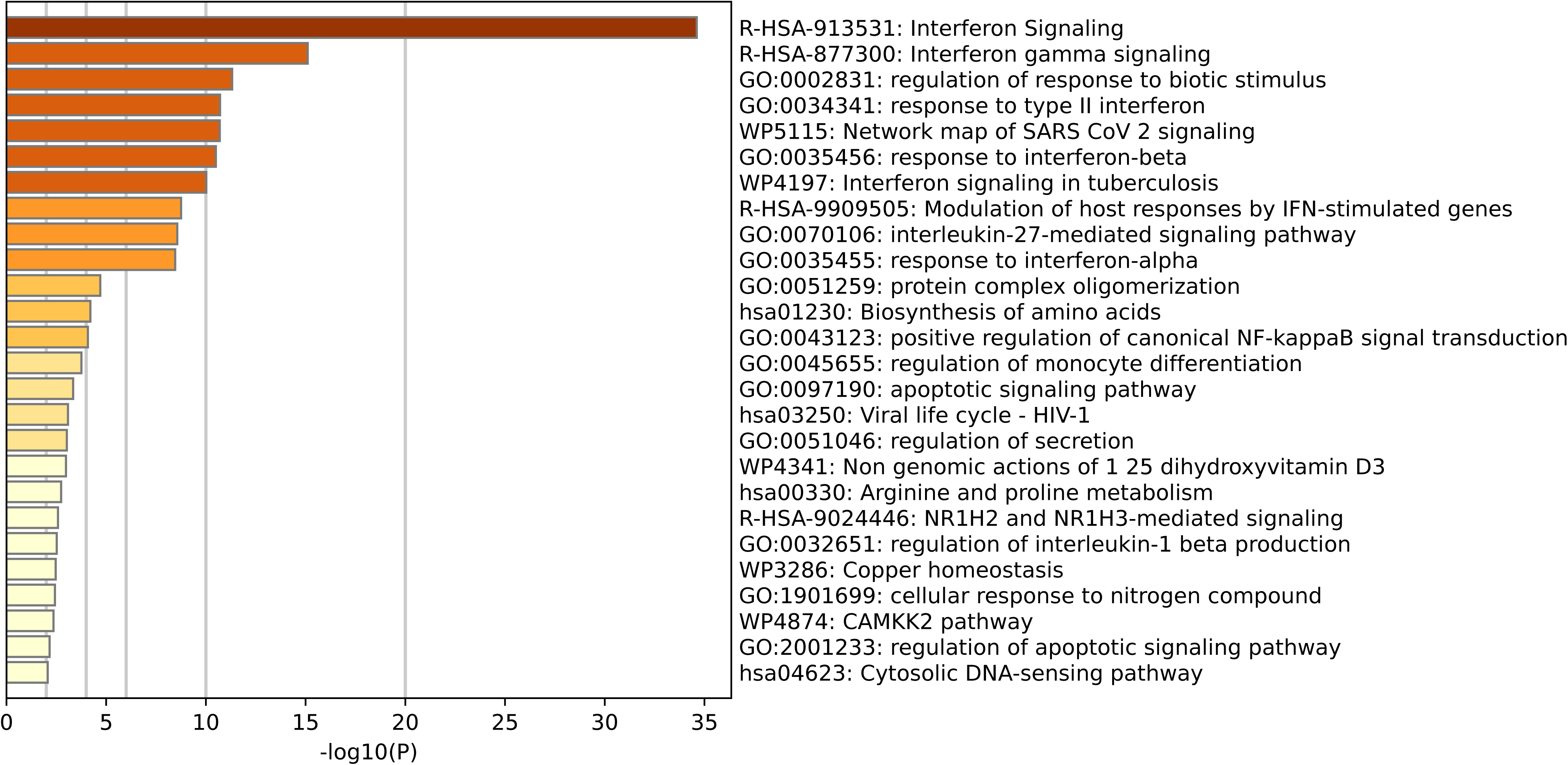
Pathway enrichment for region PE3, polychromatic erythroblasts.

**Supplementary Figure S16.**
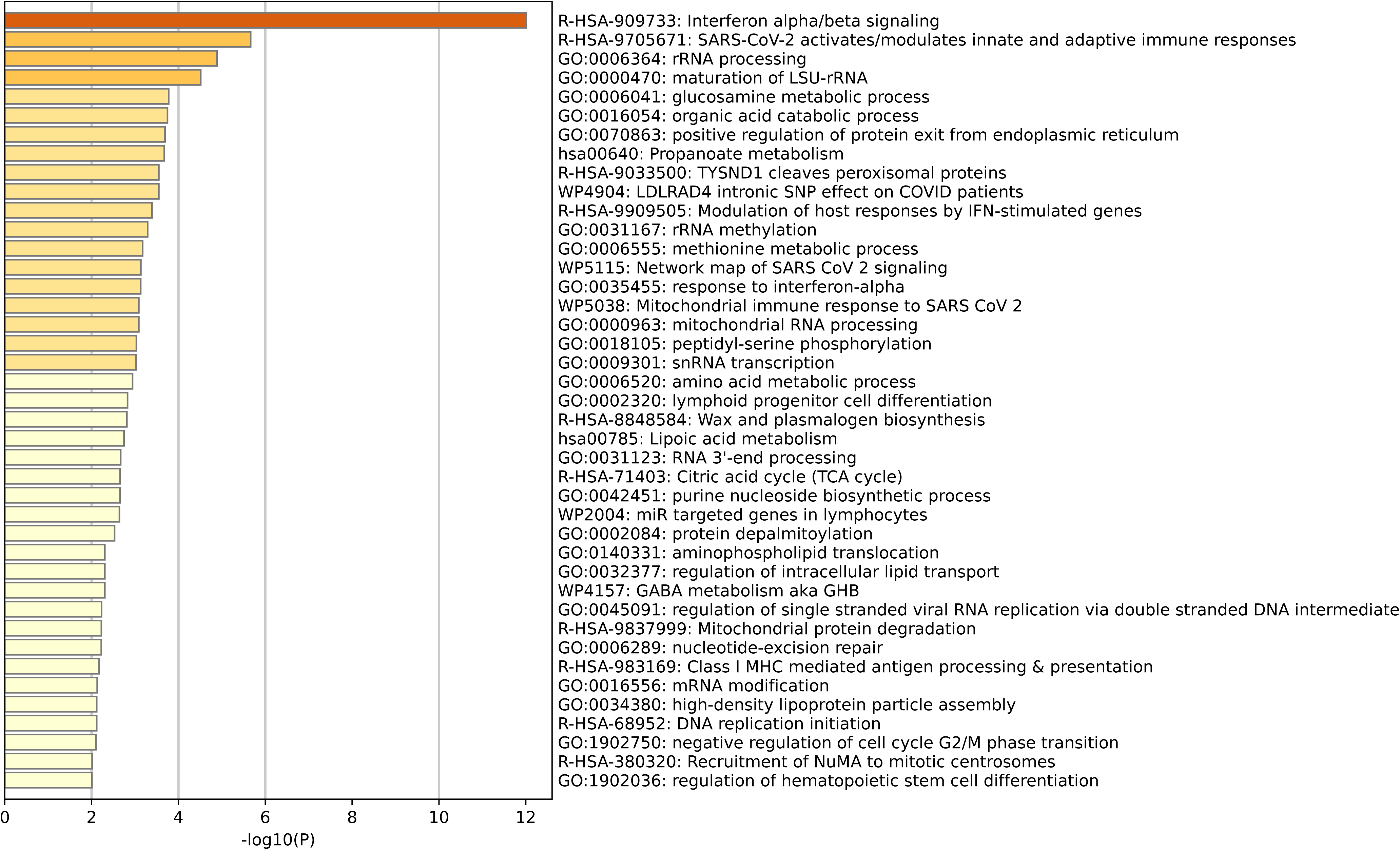
Pathway enrichment for region PE4, polychromatic erythroblasts.

**Supplementary Figure S17.**
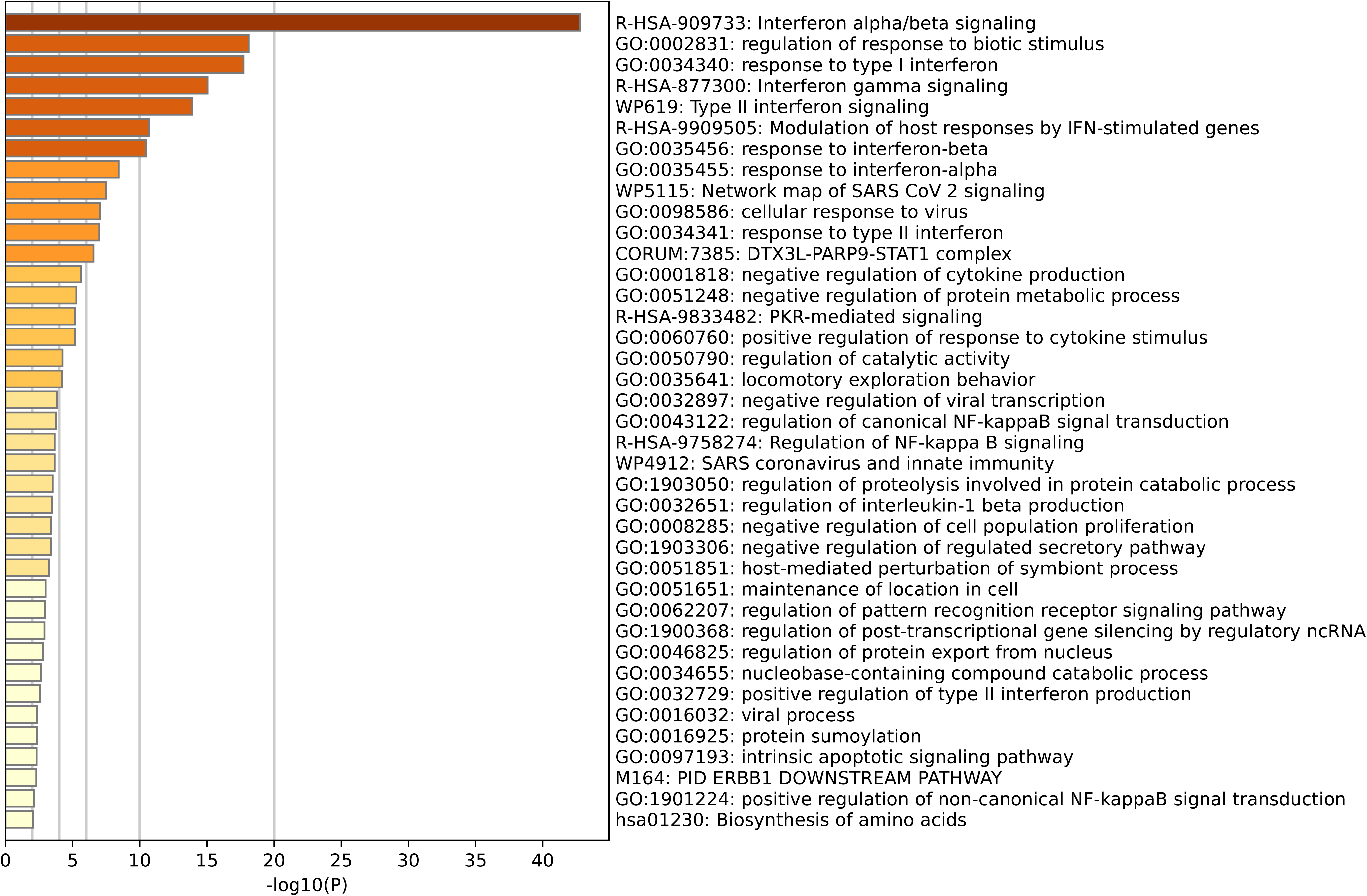
Pathway enrichment for region PE5, polychromatic erythroblasts.

**Supplementary Figure S18.**
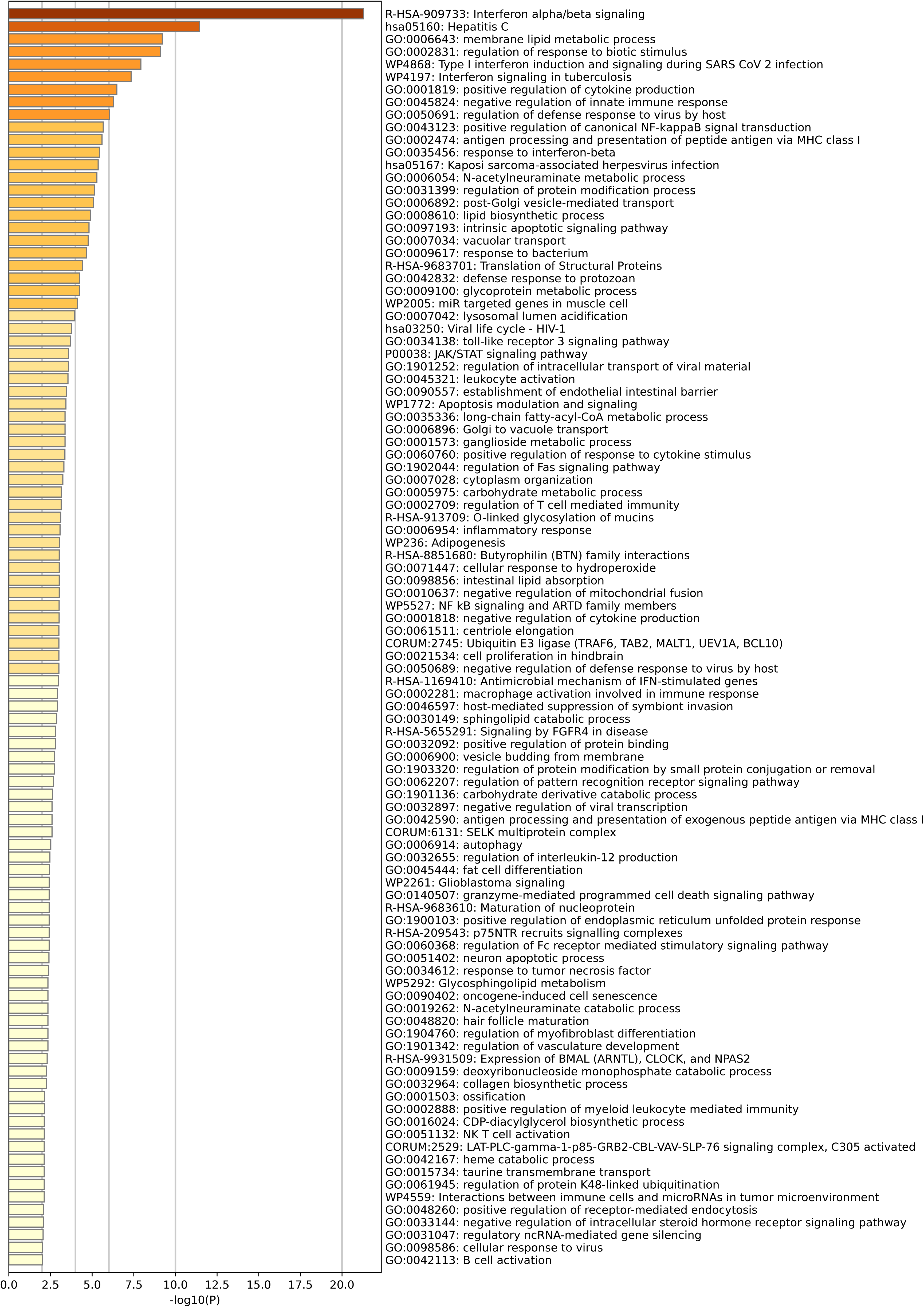
Pathway enrichment for region OE1, orthochromatic erythroblasts.

**Supplementary Figure S19.**
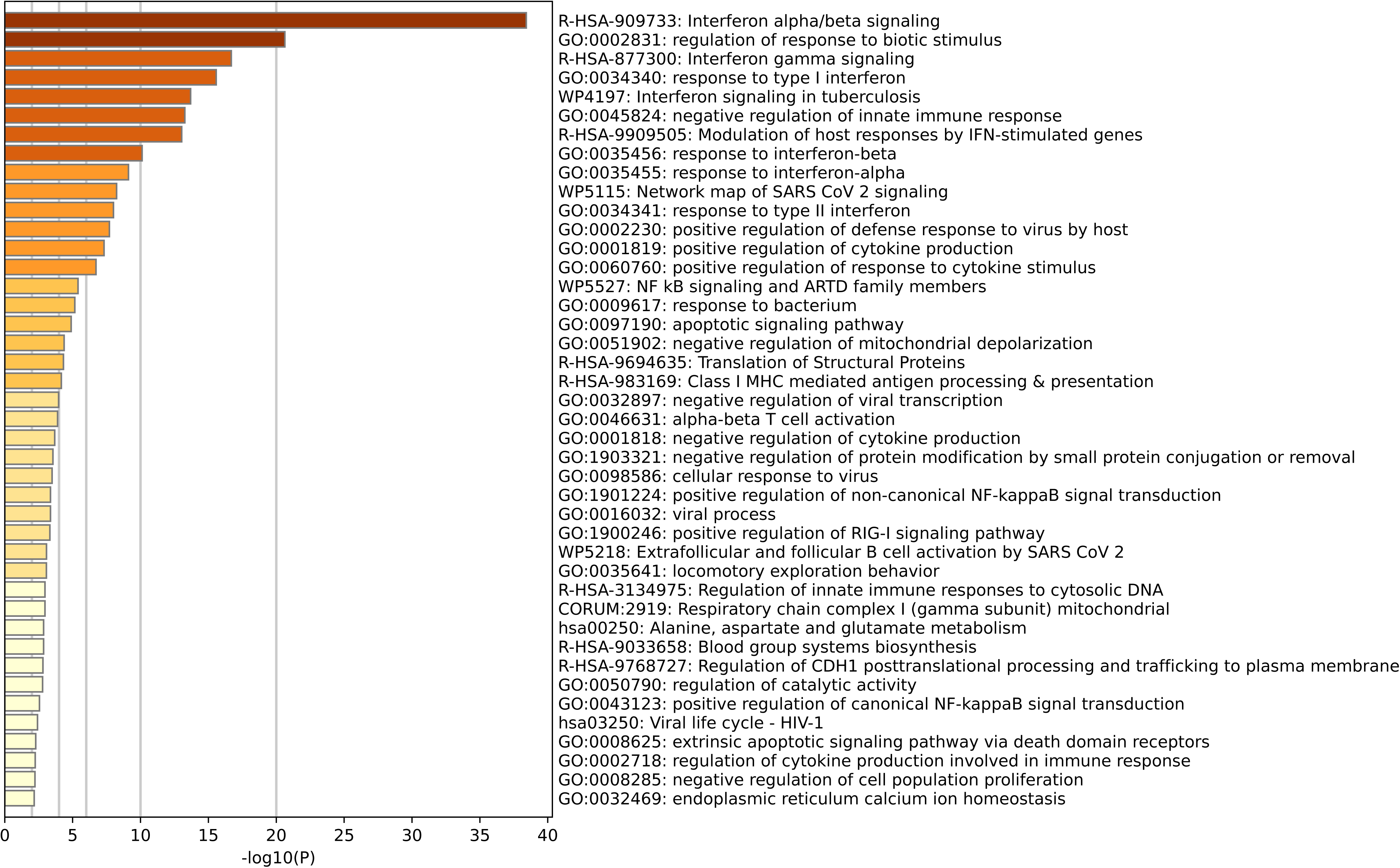
Pathway enrichment for region OE2, orthochromatic erythroblasts.

**Supplementary Figure S20.**
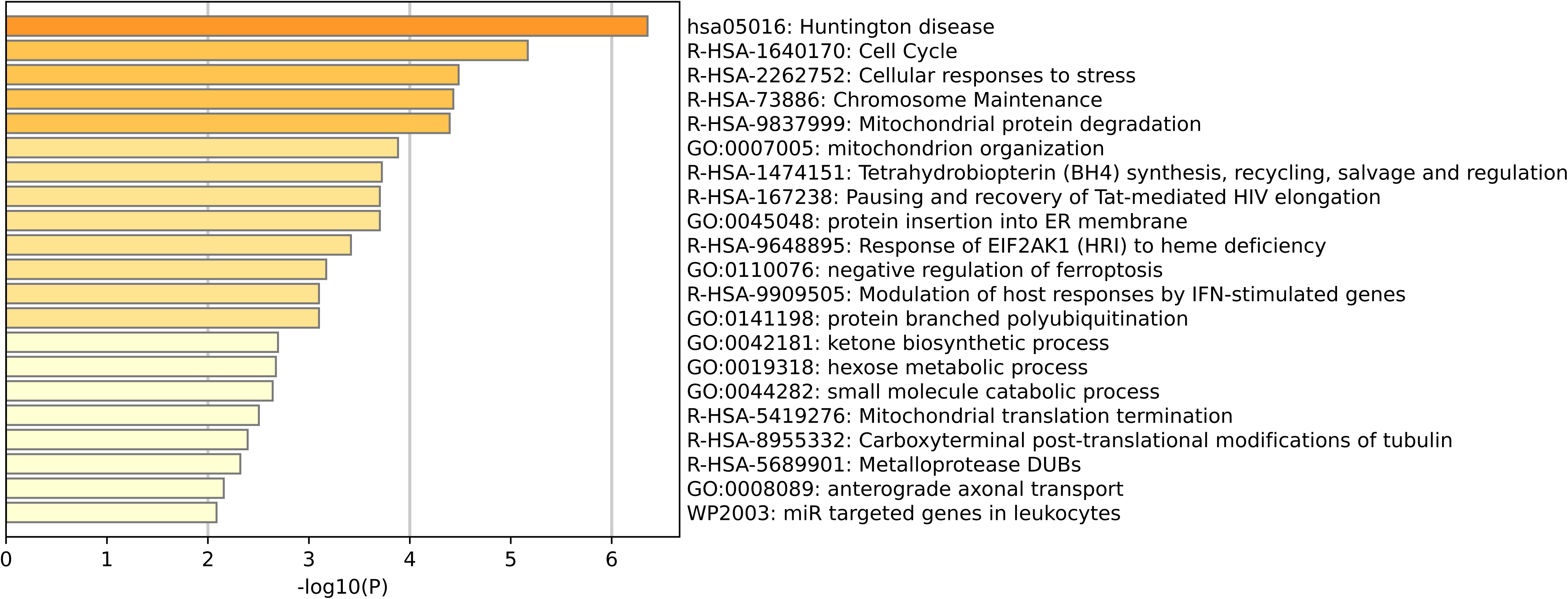
Pathway enrichment for region OE3, orthochromatic erythroblasts.

**Supplementary Figure S21.**
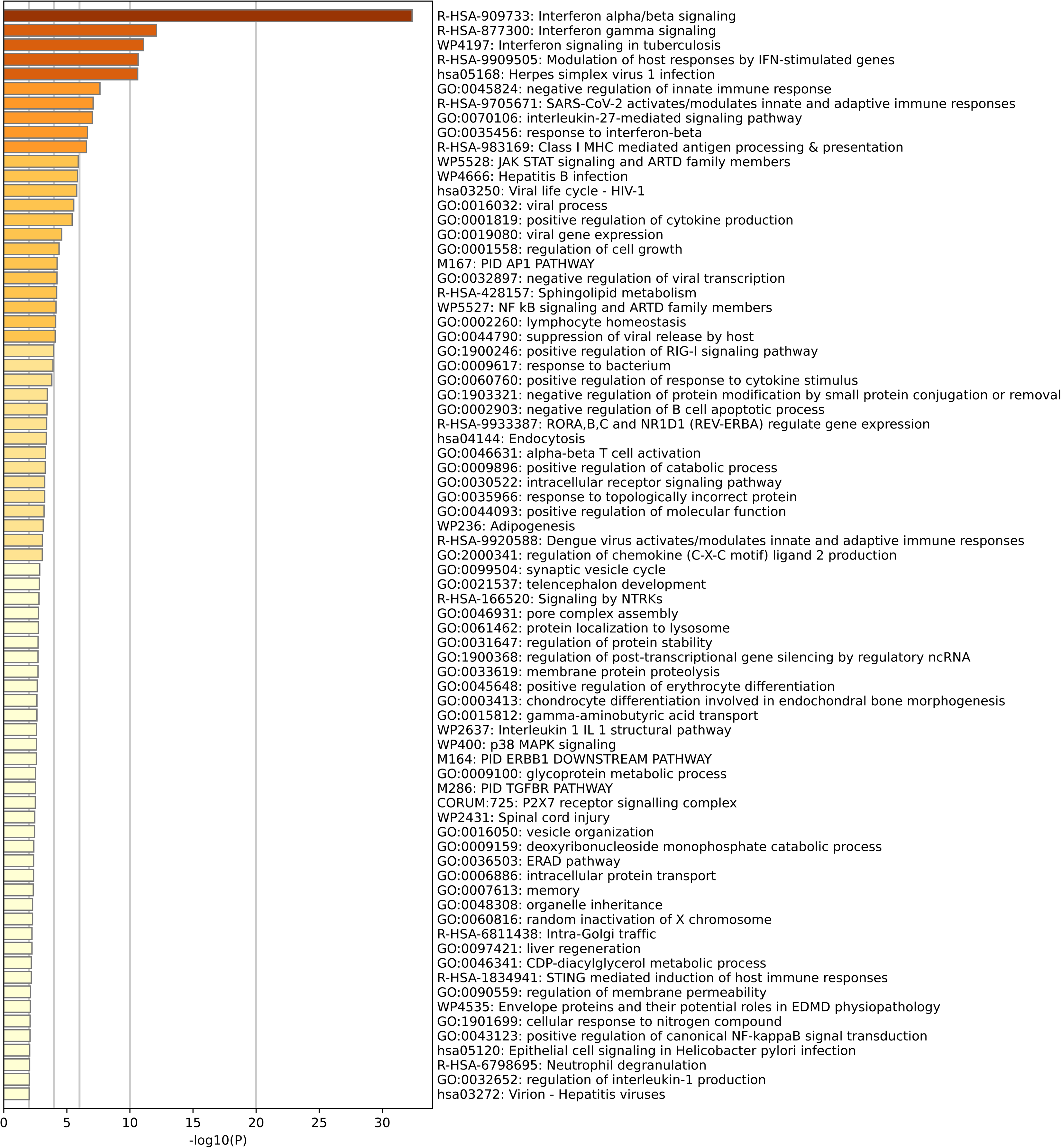
Pathway enrichment for region OE4, orthochromatic erythroblasts.

**Supplementary Figure S22.**
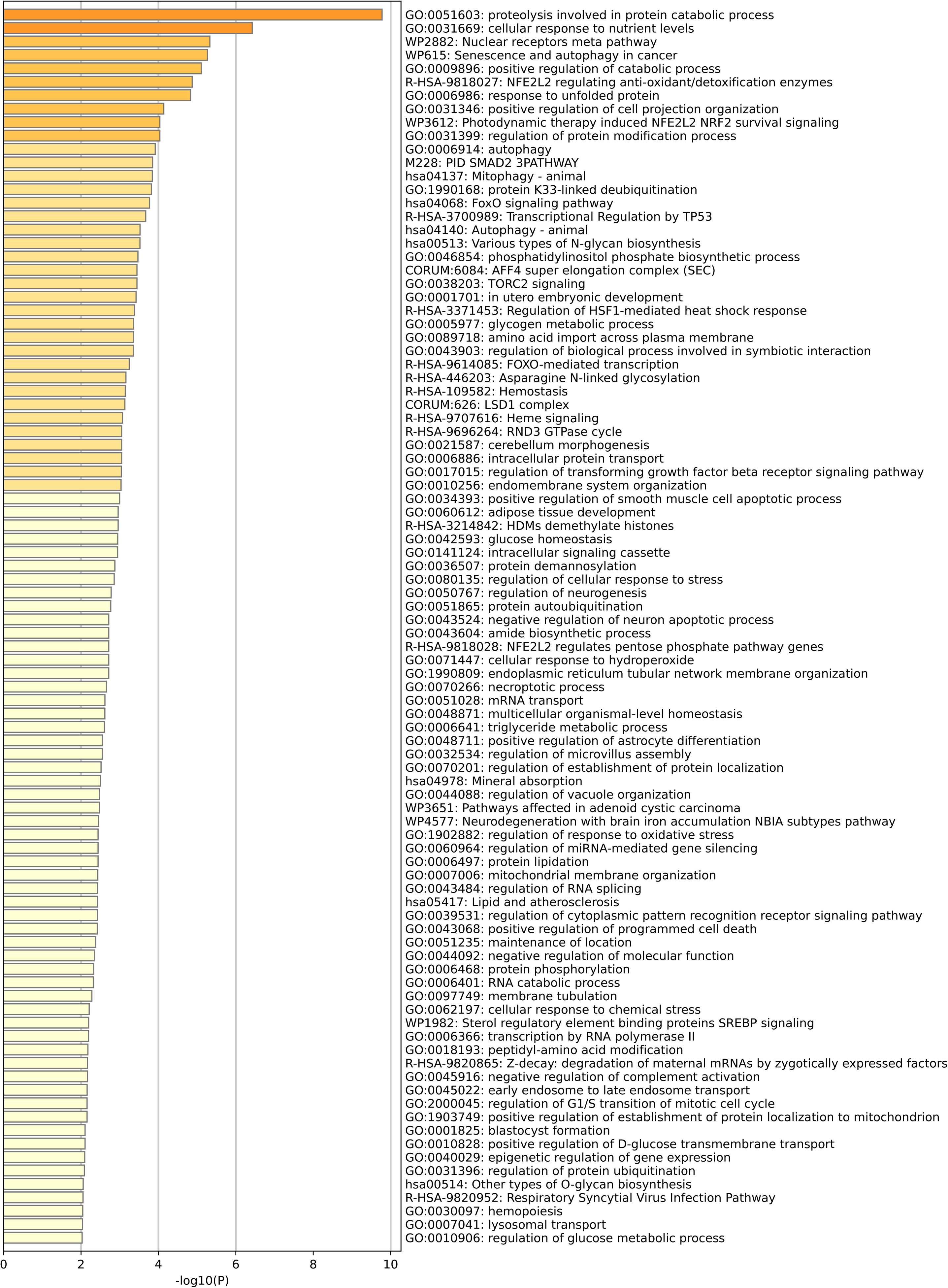
Pathway enrichment for region OE5, orthochromatic erythroblasts.

**Supplementary Figure S23.**
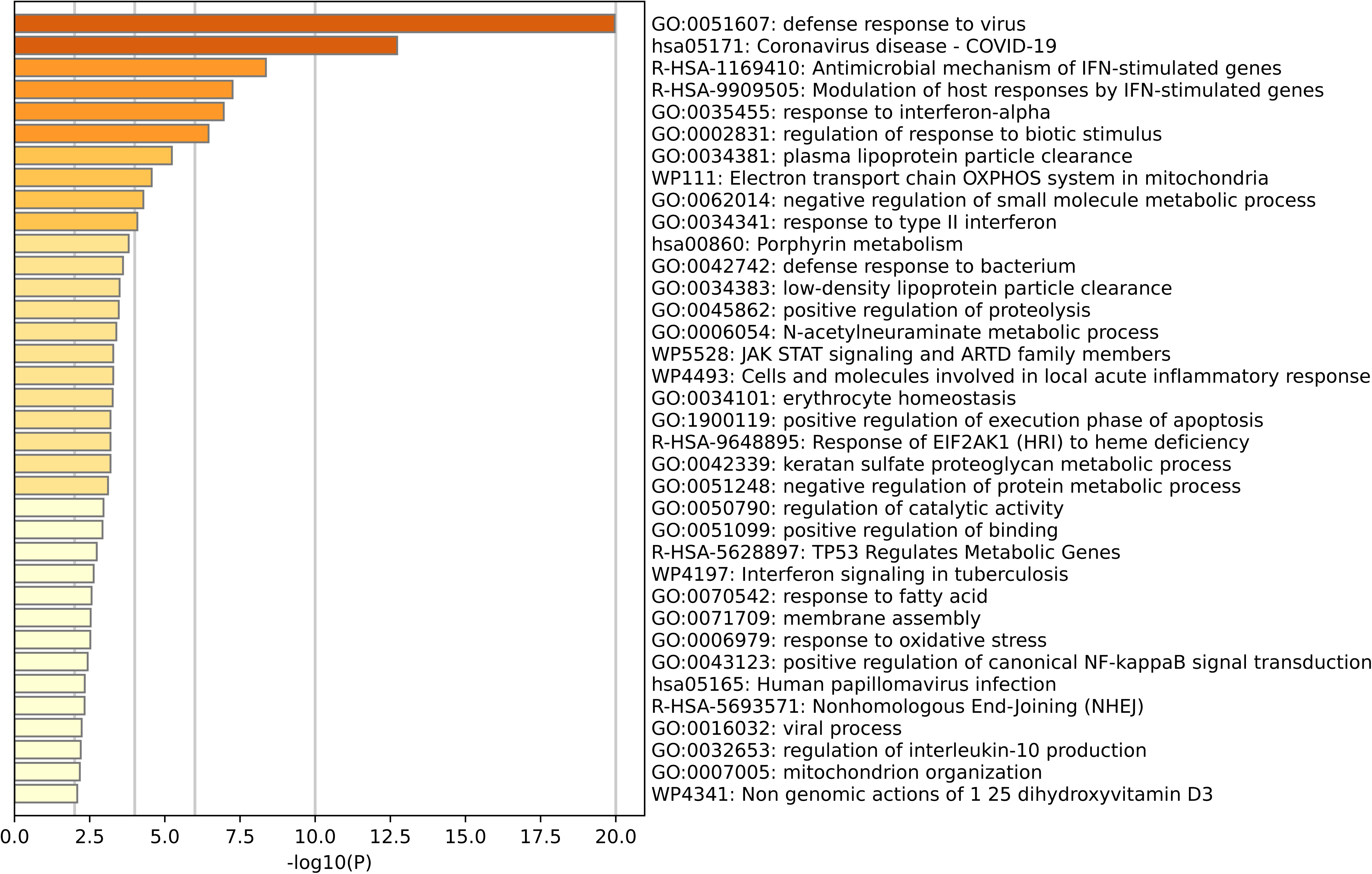
Pathway enrichment for region R1, reticulocytes.

**Supplementary Figure S24.**
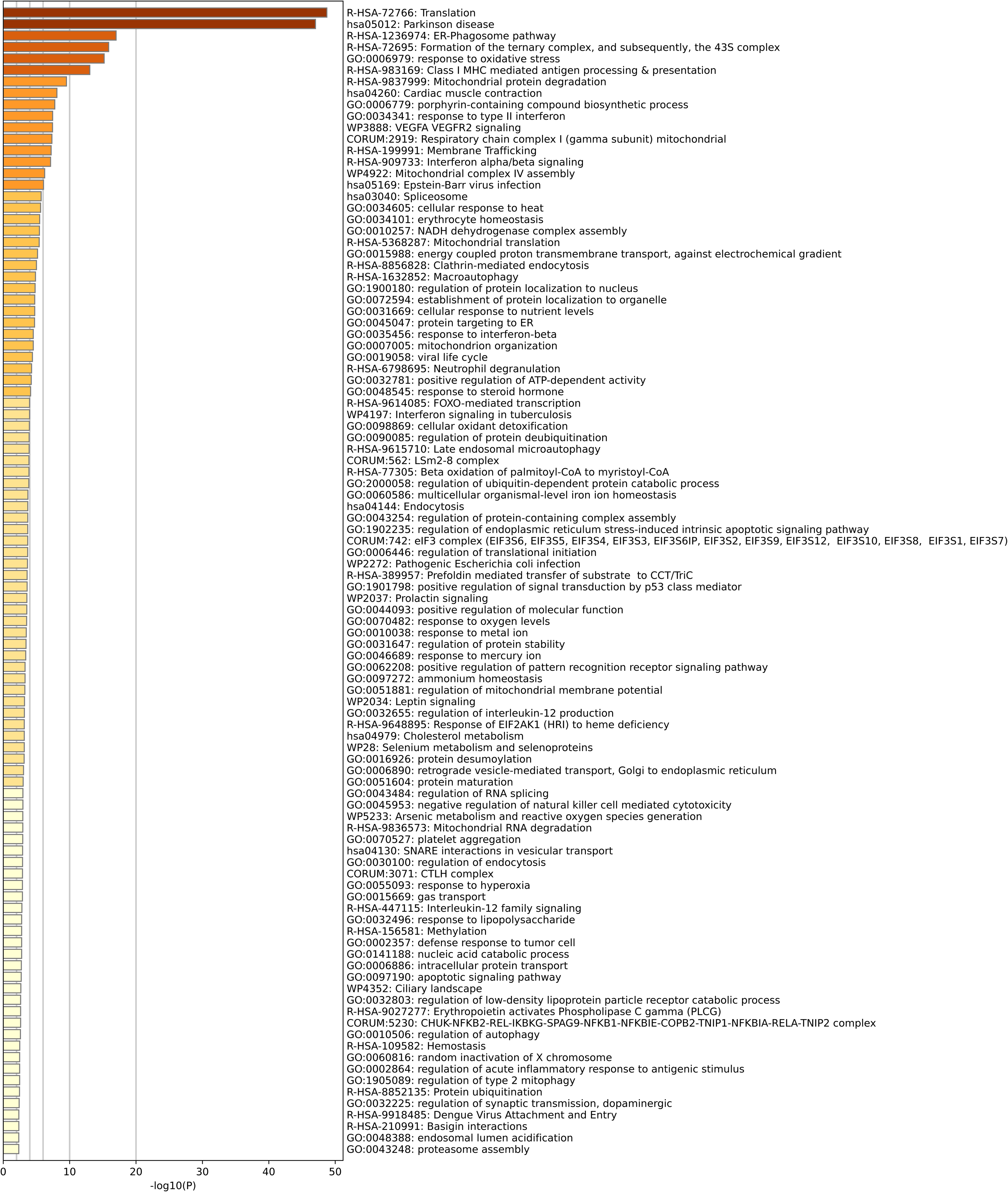
Pathway enrichment for region R2, reticulocytes.

**Supplementary Figure S25.**
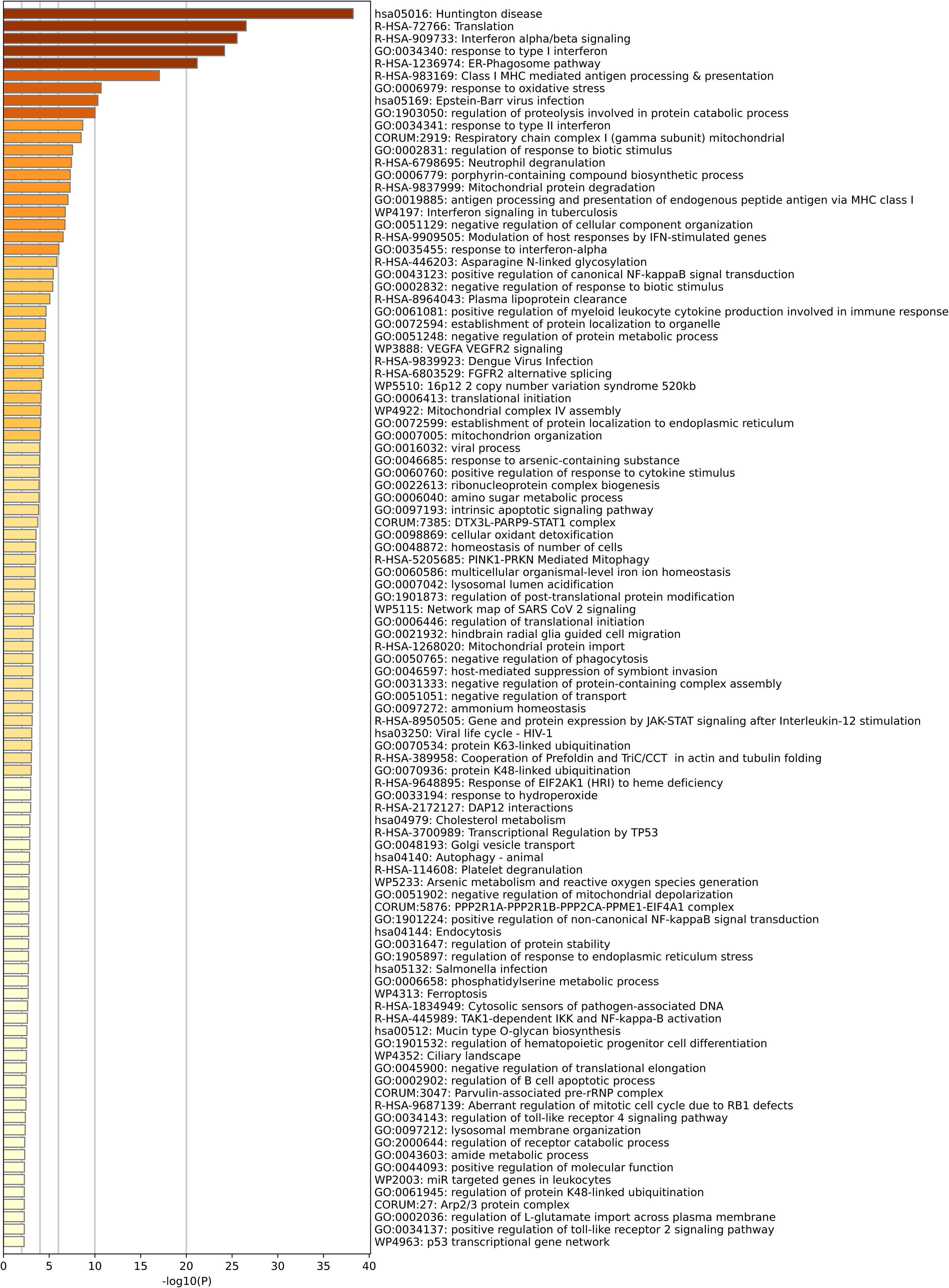
Pathway enrichment for region R3, reticulocytes.

