## Supplementary Information for "Basophilic Erythroblast Emerges as the Key Turning Point in Polycythemia Vera"

### Supplementary Methods

#### Dataset and study design

The primary dataset comprises 12 samples: 6 polycythemia vera (PV) patients and 6 healthy controls (HC). Within each disease group, 3 donors were cultured with IF-Alpha (500 U/mL IF-Alpha2b, 10 days) and 3 were left untreated, yielding four groups: Plus_PV, Minus_PV, Plus_HC, Minus_HC (Zenodo 10.5281/zenodo.14204623; Kalmer et al., 2025).

#### Preprocessing and quality control

Cells were processed with CellRanger v3.1.0, integrated and clustered with Seurat v4.2.0 by the original authors; all analyses reported in the manuscript were performed in Seurat v5.0. Cells were retained with at least 50 detected genes; genes retained if expressed in at least 5 cells. QC filtering kept cells with nFeature_RNA > 400, nCount_RNA < 40,000, and mitochondrial gene percentage below 15%. Doublets were predicted and removed with DoubletFinder using expected doublet proportions from the 10x Chromium Next GEM Single Cell 3′ v3.1 user guide (Rev. D). Samples were integrated with FindIntegrationAnchors/IntegrateData; a shared nearest-neighbour graph was built with FindNeighbors on the first 30 principal components, and clusters identified with FindClusters at resolution 0.5. An elbow plot (Supplementary Fig. S1) showed the standard deviation plateauing after the fifth principal component. Fourteen initial clusters were reduced to thirteen after removing one unidentified-cell cluster, then assigned to ten cell types: MEPs, proerythroblasts, basophilic, polychromatic and orthochromatic erythroblasts, reticulocytes, B cells, T cells, granulocytes and mast cells. No batch effect was observed across sample, stage, treatment or condition groupings (Supplementary Fig. S1).

#### Differential expression — bulk

Bulk PV data (GEO GSE26049) were analysed with the limma package (v3.54.0, R 4.2.2). Expression values were log2-transformed where indicated by a quantile check; probes with missing values were removed; probe IDs were mapped to gene symbols and Entrez IDs via hgu133plus2.db. Of the deposited GSE26049 samples, 41 PV and 21 control samples were used (essential thrombocythemia and primary myelofibrosis samples excluded). A gene was called differentially expressed at $|\log2FC| > 1.0$ and adjusted $P < 0.05$. Functional enrichment of the resulting up- and downregulated gene sets was performed with Metascape; these results are shown as panels a and b of main-text Figure 3.

#### Differential expression — single cell

scRNA-seq DEGs were identified with Seurat's FindMarkers() function using the Wilcoxon rank-sum test, first by pooling all cells per condition (disregarding cell type) and then repeated within each of the 10 annotated cell types, across seven pairwise comparisons: PV vs HC, Minus_PV vs Minus_HC, Plus_PV vs Plus_HC, Plus vs Minus, Plus_HC vs Minus_HC, Plus_PV vs Minus_PV, and the bulk PV vs HC comparison used for validation. The significance criteria matched the bulk analysis except for a lower average log2FC threshold of 0.5 (adjusted $P < 0.05$ in both cases). Disease-effect genes were defined as those differentially expressed in every PV vs HC comparison; treatment-effect genes as those differentially expressed in every Plus vs Minus comparison. Requiring overlap across all relevant comparisons, cross-validated against the bulk DEGs, reduced false positives (Supplementary Table S2).

#### Enrichment analysis

GO and KEGG enrichment used Metascape, which merges near-duplicate terms into a single representative term (lowest adjusted P value retained) and draws on GO, KEGG, DisGeNET, TRRUST, WikiPathways and Reactome, among other sources. JAK-STAT pathway visualization used Pathview (hsa04630).

#### NicheNet analysis

NicheNet was run with the latest ligand-target matrix, ligand-receptor network and weighted integrated networks downloaded from their respective Zenodo repositories, using default parameters. Six pairwise comparisons (PV vs HC, Minus_PV vs Minus_HC, Plus_PV vs Plus_HC, Plus_PV vs Minus_PV, Plus_HC vs Minus_HC, Plus vs Minus) were performed for every cell type as receiver.

#### Pseudotime trajectory analysis

Pseudotime was computed per cell type with Monocle3's order_cells(), which projects each cell onto a principal graph learned from the UMAP embedding via reversed graph embedding (SimplePPT), assigning pseudotime as the geodesic distance from the cell's projected position to a user-defined root node representing the most immature progenitor population. Comparison across the four condition-stratified trajectories (Minus_HC, Minus_PV, Plus_HC, Plus_PV) revealed condition-dependent structure specifically in Basophilic Erythroblasts, Polychromatic Erythroblasts, Orthochromatic Erythroblasts, Reticulocytes and T cells; MEPs, Granulocytes, Mast cells and B cells showed none. For cell types with observed structure, representative regions at the trajectory extremes were selected (excluding transitional/intermediate cells) for region-based differential expression and enrichment: 6 regions in Basophilic Erythroblasts (BE1–BE6), 5 in Polychromatic Erythroblasts (PE1–PE5), 5 in Orthochromatic Erythroblasts (OE1–OE5), 3 in Reticulocytes (R1–R3), and 6 clusters (3 main, 2 sub-areas each) in T cells, though the T cell trajectory characteristics were judged too uncertain for further analysis (Supplementary Table S3 and Supplementary Table S4; Supplementary Figs. S5 and S6; region enrichment figures in the companion file Supplementary_Figures_S7-S25_Enrichment.pdf).

#### InferCNV analysis

Copy number variation was inferred with inferCNV using a raw count matrix and a gene-order list (chromosome, start, end). A gene-filtering cutoff of 0.1 was applied (recommended for 10x Genomics data). No reference cells were supplied for whole-object runs (package uses the mean of all cells as baseline); for cell-type-specific runs, B cells, T cells, granulocytes and mast cells served as the reference population. The mean absolute CNV score threshold was defined as the maximum of two estimators computed from the reference-cell CNV score distribution: the median plus three median absolute deviations (MAD), and the 99th percentile; the resulting threshold was 0.02570775. The percentage of cells exceeding this threshold was then computed for each erythroid stage; these are the values reported in the manuscript (Supplementary Table S5; Supplementary Fig. S26). Genes correlated with the per-cell CNV score were identified by Spearman correlation across all cells, ranked by absolute correlation coefficient, and retained above a mean absolute correlation of 0.13 (Supplementary Table S5).

#### Drug repurposing analysis

Disease-affected genes identified at the basophilic erythroblast stage were considered against known and investigational agents relevant to PV and other myeloproliferative neoplasms, evaluated by mechanism of action, clinical development stage, and existing MPN evidence (Supplementary Table S7).

#### Statistics and reproducibility

Consolidated significance thresholds used throughout this study: bulk DEGs, |log2FC| > 1.0 and adjusted P < 0.05 (limma); single-cell DEGs, |log2FC| > 0.5 and adjusted P < 0.05, Wilcoxon rank-sum test (Seurat FindMarkers); CNV-correlated genes, mean absolute Spearman correlation > 0.13.

### Supplementary Figures

#### Supplementary Figure S1. Quality control and batch-effect assessment.

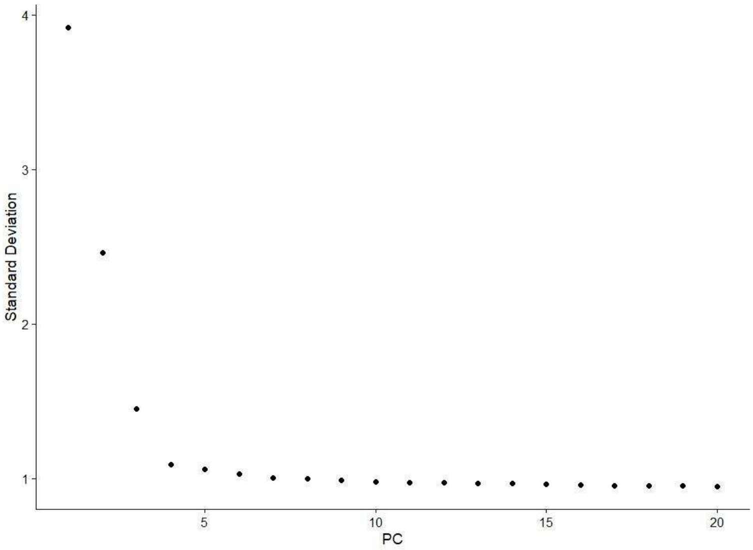


(a) Principal component elbow plot; standard deviation plateaus at approximately 1.0 after the fifth principal component.

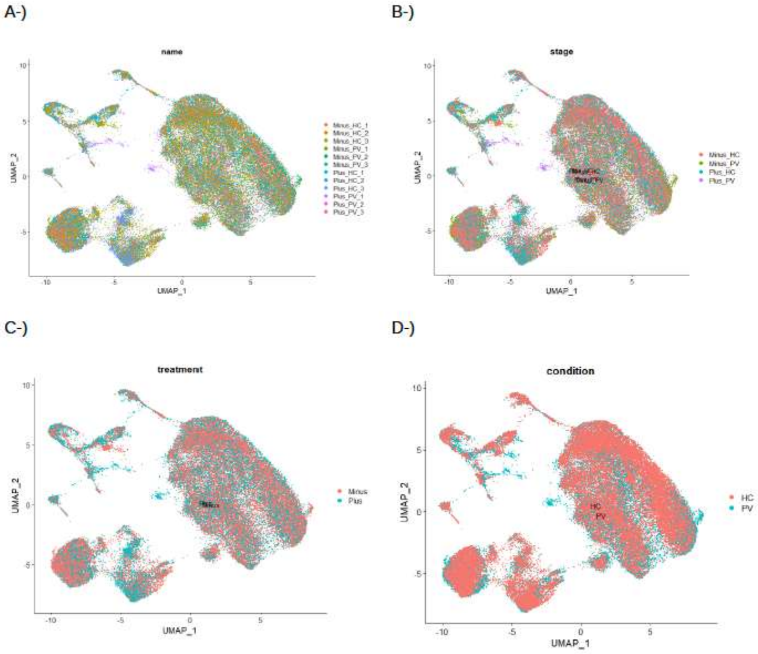

(b) UMAP embeddings coloured by sample name, stage, treatment and condition; no systematic batch structure is apparent in any grouping.

**Supplementary Figure S2. NicheNet ligand-receptor analysis for polychromatic erythroblasts.**

(a) Prior interaction potentials between prioritised ligands and their receptors.

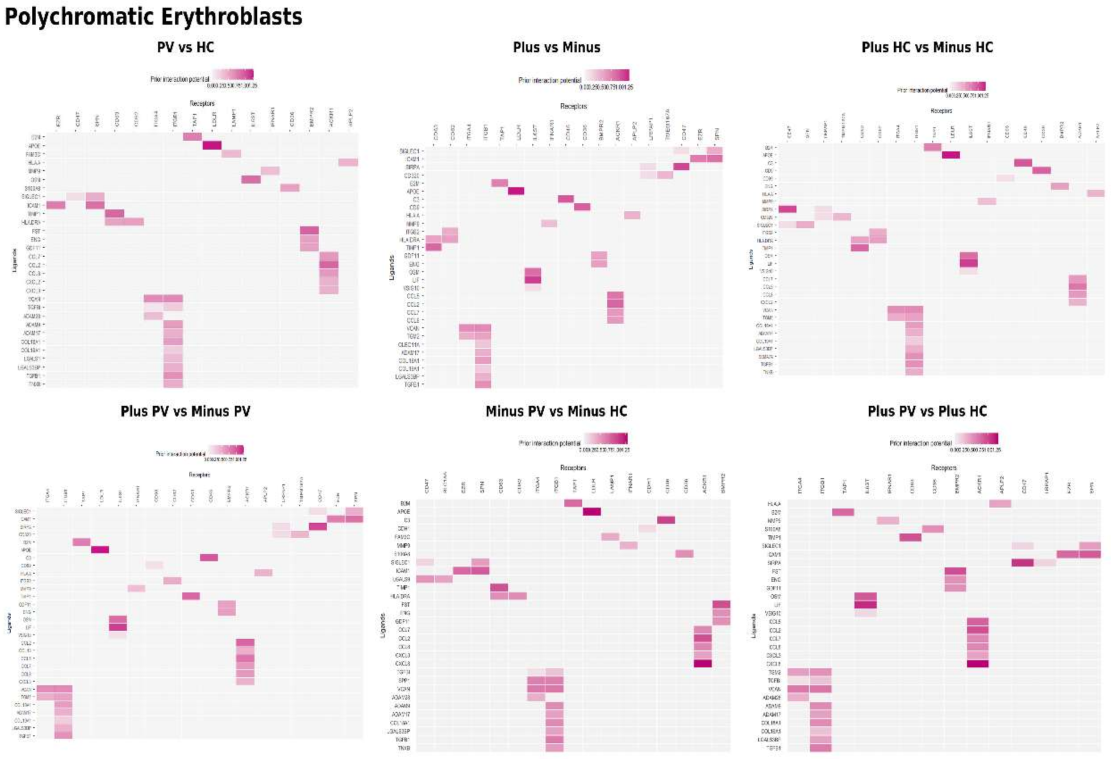


(b) Ligand expression log fold change in the sender populations.

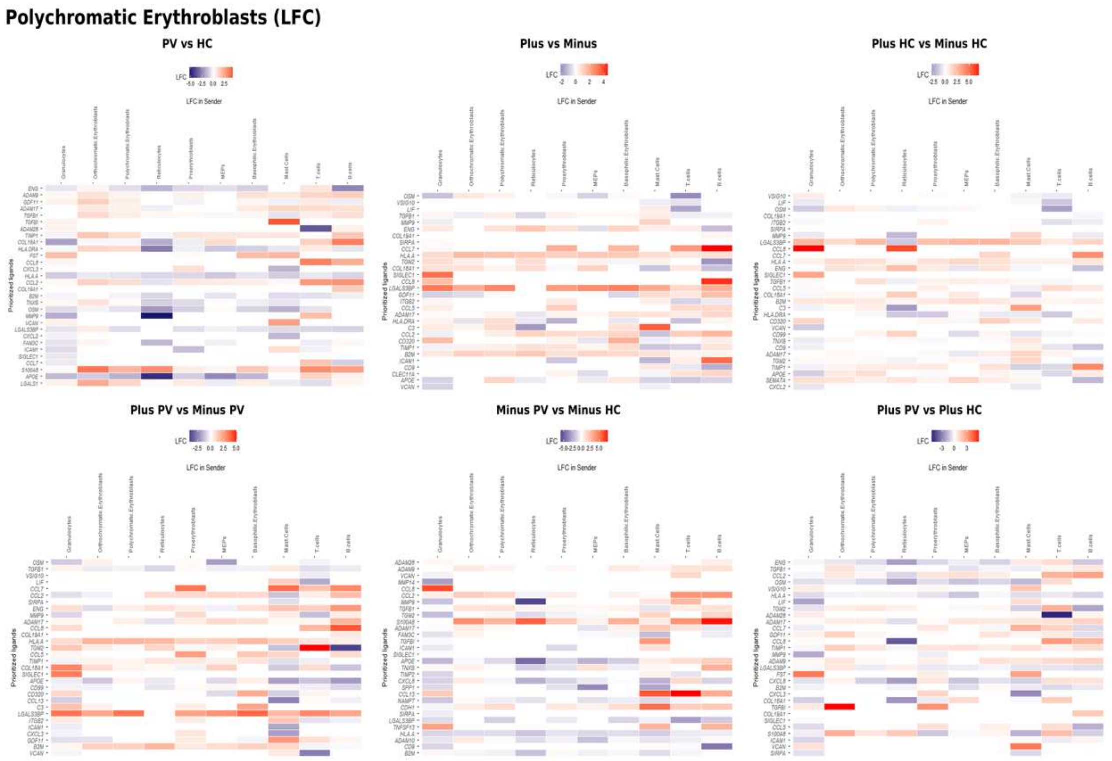


Panels correspond to the six pairwise comparisons: PV versus HC, Plus versus Minus, Plus_HC versus Minus_HC, Plus_PV versus Minus_PV, Minus_PV versus Minus_HC, and Plus_PV versus Plus_HC. At this stage the disease comparisons add BMP9, LGALS1, LGALS3BP, VCAN, ADAM28, TGFBI and TGM2 to the erythroid repertoire, engaging CD47, SPN, ITGB1, TAP1 and IFNAR1.

### Supplementary Figure S3. NicheNet ligand-receptor analysis for orthochromatic erythroblasts.

(a) Prior interaction potentials between prioritised ligands and their receptors.

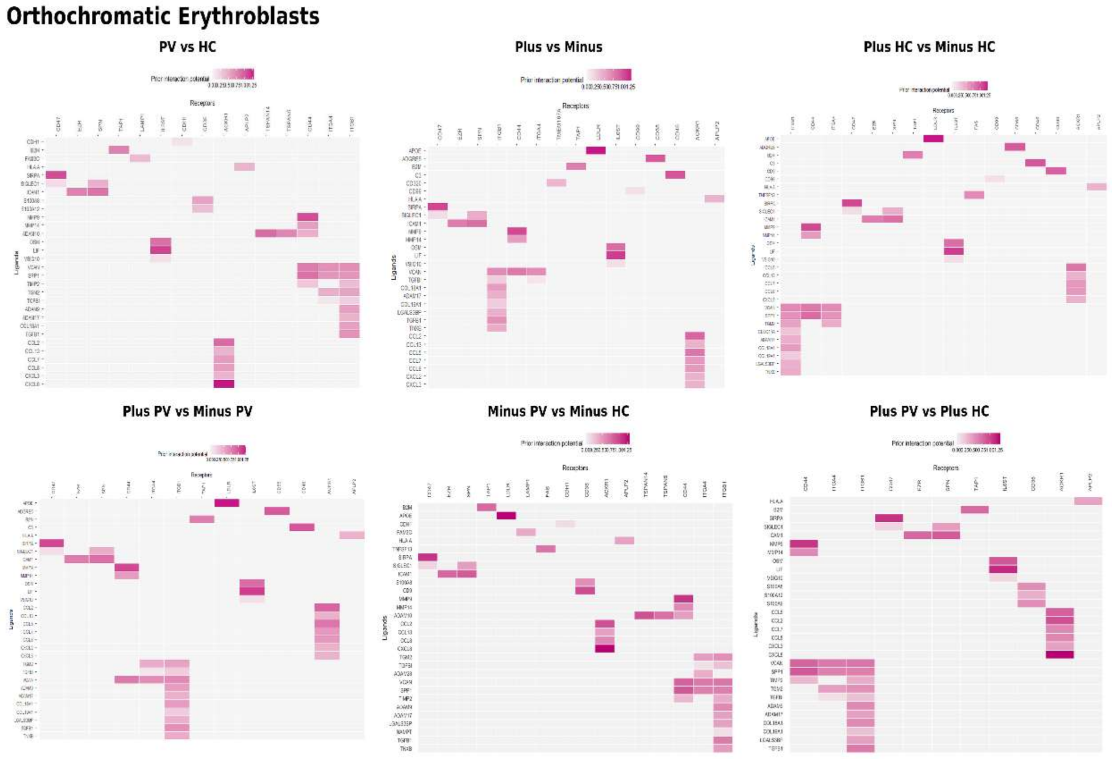

(b) Ligand expression log fold change in the sender populations.

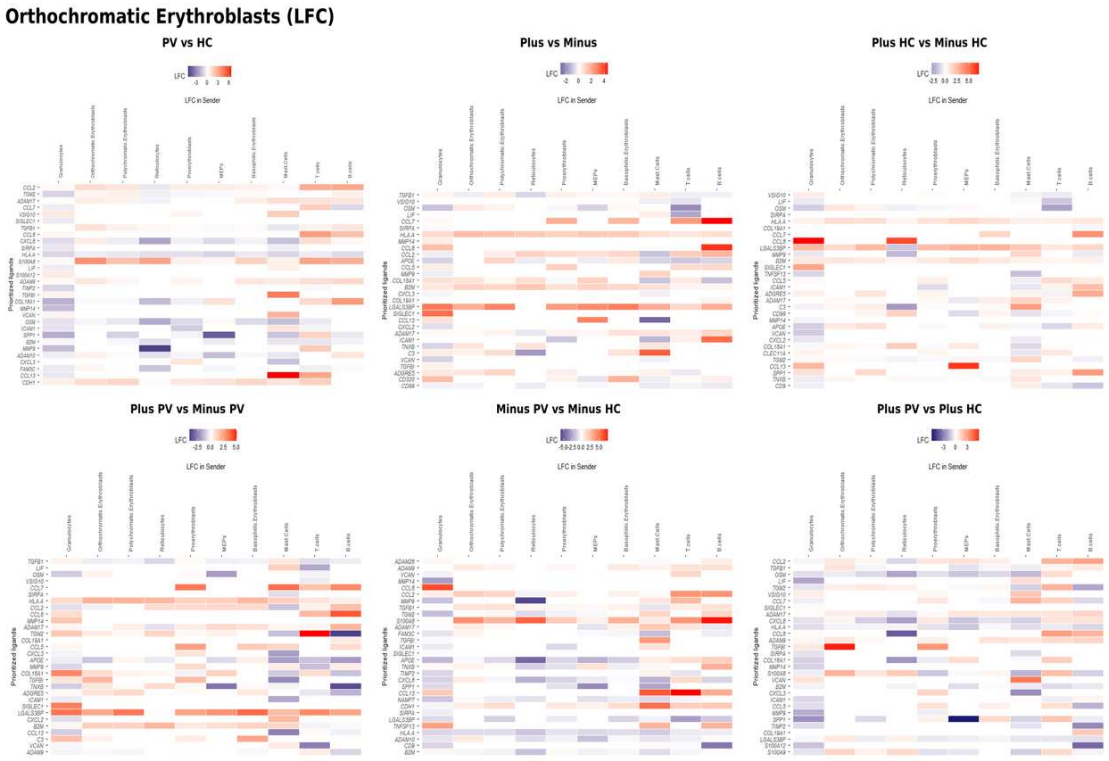


Panels correspond to the six pairwise comparisons: PV versus HC, Plus versus Minus, Plus_HC versus Minus_HC, Plus_PV versus Minus_PV, Minus_PV versus Minus_HC, and Plus_PV versus Plus_HC. The SIRPA, SIGLEC1 and S100A axes established at earlier stages are reinforced, and CCL2, ADAM17, SIGLEC1, TGM2 and VCAN are broadly upregulated in the disease comparisons.

### Supplementary Figure S4. NicheNet ligand-receptor analysis for reticulocytes.

(a) Prior interaction potentials between prioritised ligands and their receptors.

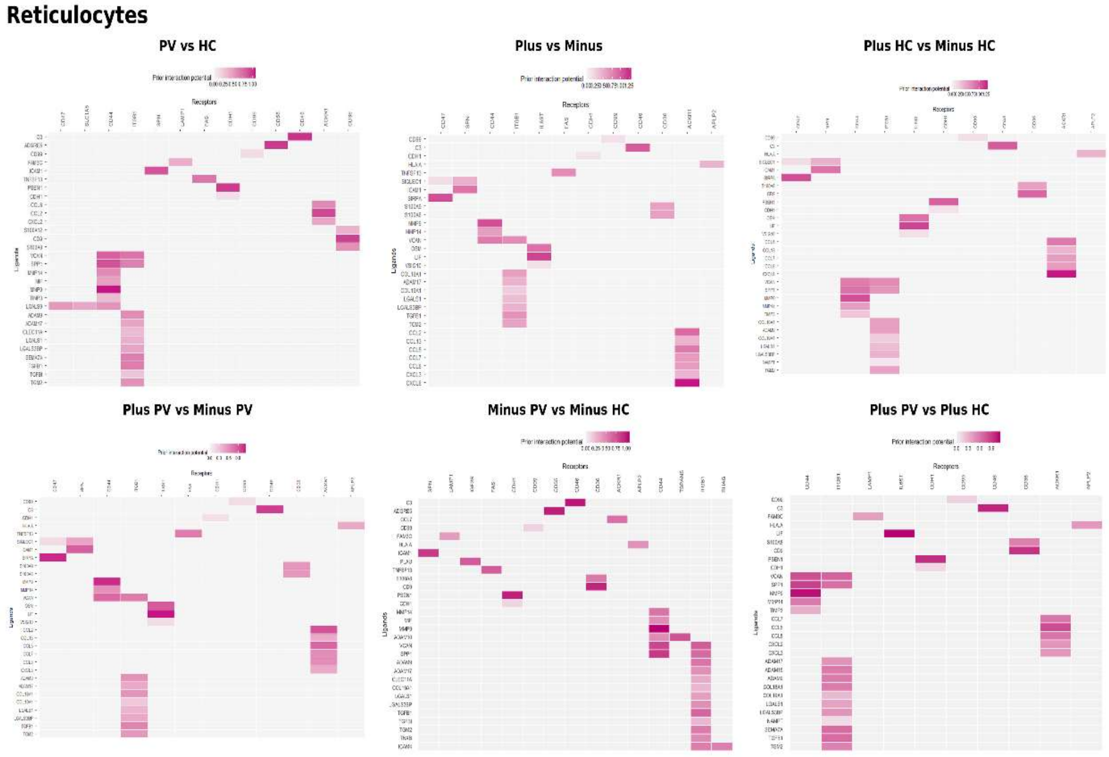

(b) Ligand expression log fold change in the sender populations.

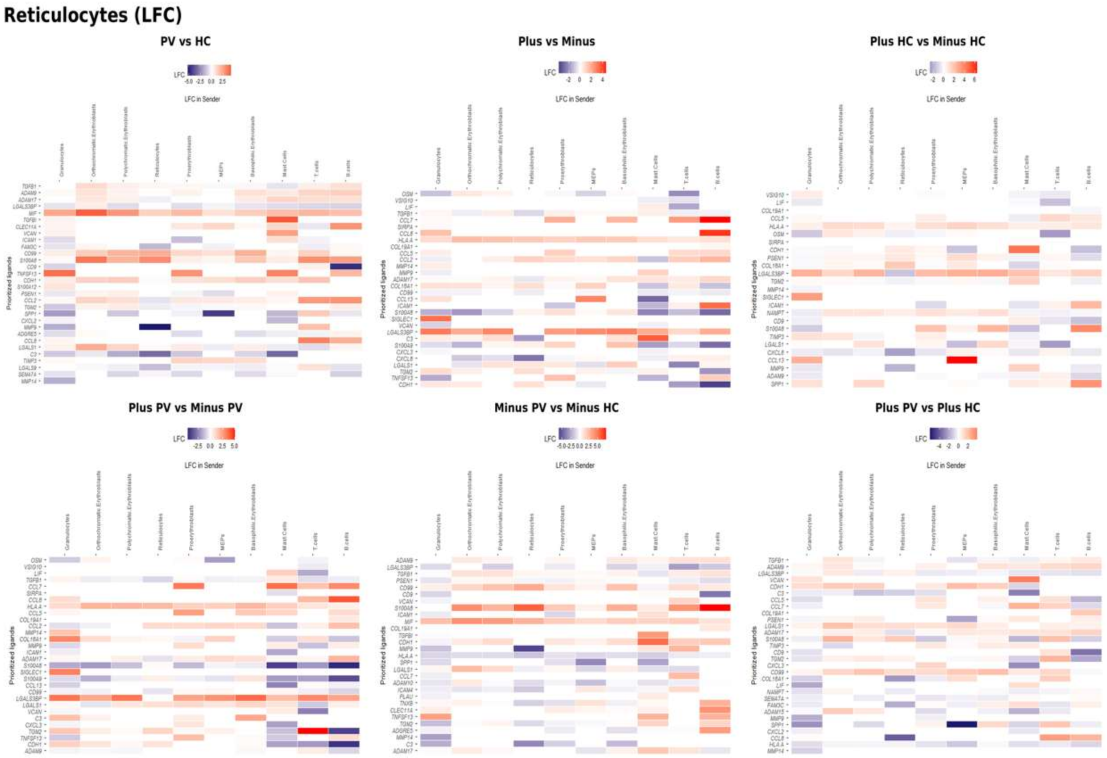


Panels correspond to the six pairwise comparisons: PV versus HC, Plus versus Minus, Plus_HC versus Minus_HC, Plus_PV versus Minus_PV, Minus_PV versus Minus_HC, and Plus_PV versus Plus_HC. CD99-PILRA and PSEN1 interactions are relatively specific to this terminal stage, while TGFBI, ADAM9 and LGALS3BP remain upregulated in the disease comparisons.

**Supplementary Figure S5. Pseudotime trajectory of megakaryocyte-erythroid progenitors.**

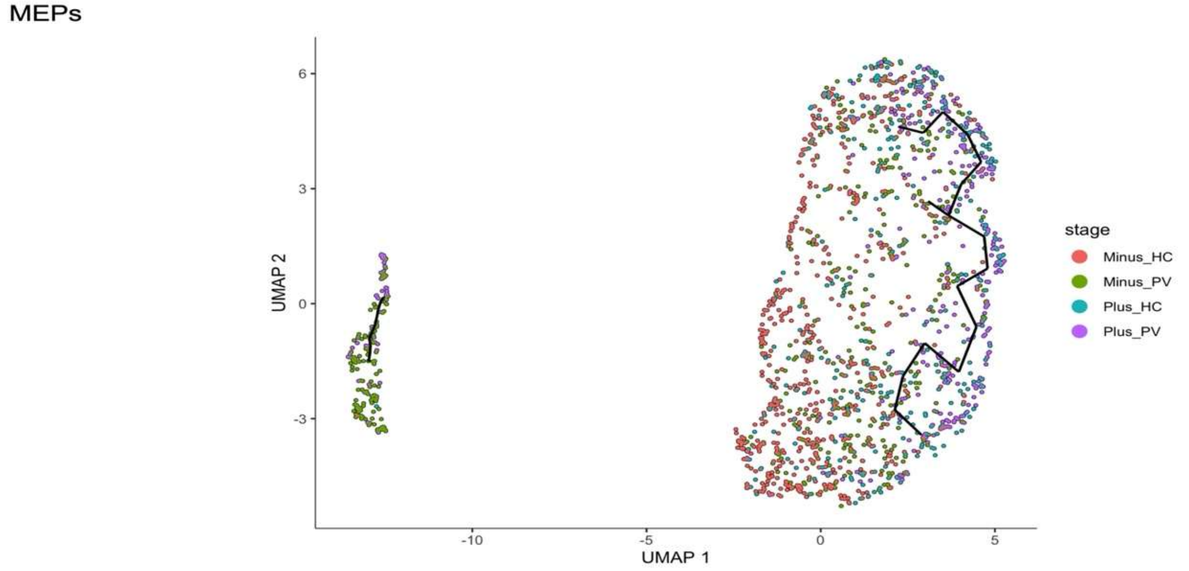


Monocle3 trajectory for MEPs, coloured by condition (Minus_HC, Minus_PV, Plus_HC, Plus_PV). No condition-dependent structure is observed, consistent with the disease effect not yet being detectable at this earliest erythroid progenitor stage.

**Supplementary Figure S6. Pseudotime trajectory of proerythroblasts.**

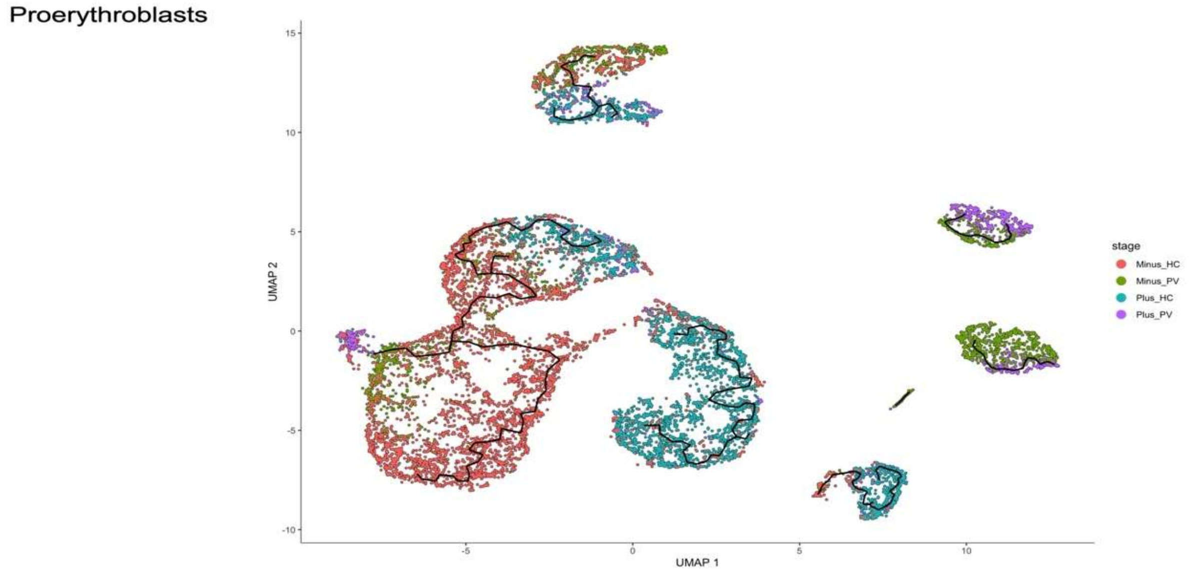


Monocle3 trajectory for proerythroblasts, coloured by condition. Cells from all four conditions are intermixed along the trajectory, with no condition-dependent structure, despite this stage already carrying a substantial disease-effect gene signature.

**Supplementary Figure S7.** Pathway enrichment for region BE1 (basophilic erythroblasts). Metascape enrichment of the genes differentially expressed in BE1 relative to all remaining cells of the same cell type; bars show -log10(P).

**Supplementary Figure S8.** Pathway enrichment for region BE2 (basophilic erythroblasts). Metascape enrichment of the genes differentially expressed in BE2 relative to all remaining cells of the same cell type; bars show -log10(P).

**Supplementary Figure S9.** Pathway enrichment for region BE3 (basophilic erythroblasts). Metascape enrichment of the genes differentially expressed in BE3 relative to all remaining cells of the same cell type; bars show -log10(P).

**Supplementary Figure S10.** Pathway enrichment for region BE4 (basophilic erythroblasts). Metascape enrichment of the genes differentially expressed in BE4 relative to all remaining cells of the same cell type; bars show -log10(P).

**Supplementary Figure S11.** Pathway enrichment for region BE5 (basophilic erythroblasts). Metascape enrichment of the genes differentially expressed in BE5 relative to all remaining cells of the same cell type; bars show -log10(P).

**Supplementary Figure S12.** Pathway enrichment for region BE6 (basophilic erythroblasts). Metascape enrichment of the genes differentially expressed in BE6 relative to all remaining cells of the same cell type; bars show -log10(P).

**Supplementary Figure S13.** Pathway enrichment for region PE1 (polychromatic erythroblasts). Metascape enrichment of the genes differentially expressed in PE1 relative to all remaining cells of the same cell type; bars show -log10(P).

**Supplementary Figure S14.** Pathway enrichment for region PE2 (polychromatic erythroblasts). Metascape enrichment of the genes differentially expressed in PE2 relative to all remaining cells of the same cell type; bars show -log10(P).

**Supplementary Figure S15.** Pathway enrichment for region PE3 (polychromatic erythroblasts). Metascape enrichment of the genes differentially expressed in PE3 relative to all remaining cells of the same cell type; bars show -log10(P).

**Supplementary Figure S16.** Pathway enrichment for region PE4 (polychromatic erythroblasts). Metascape enrichment of the genes differentially expressed in PE4 relative to all remaining cells of the same cell type; bars show -log10(P).

**Supplementary Figure S17.** Pathway enrichment for region PE5 (polychromatic erythroblasts). Metascape enrichment of the genes differentially expressed in PE5 relative to all remaining cells of the same cell type; bars show -log10(P).

**Supplementary Figure S18.** Pathway enrichment for region OE1 (orthochromatic erythroblasts). Metascape enrichment of the genes differentially expressed in OE1 relative to all remaining cells of the same cell type; bars show -log10(P).

**Supplementary Figure S19.** Pathway enrichment for region OE2 (orthochromatic erythroblasts). Metascape enrichment of the genes differentially expressed in OE2 relative to all remaining cells of the same cell type; bars show -log10(P).

**Supplementary Figure S20.** Pathway enrichment for region OE3 (orthochromatic erythroblasts). Metascape enrichment of the genes differentially expressed in OE3 relative to all remaining cells of the same cell type; bars show -log10(P).

**Supplementary Figure S21.** Pathway enrichment for region OE4 (orthochromatic erythroblasts). Metascape enrichment of the genes differentially expressed in OE4 relative to all remaining cells of the same cell type; bars show -log10(P).

**Supplementary Figure S22.** Pathway enrichment for region OE5 (orthochromatic erythroblasts). Metascape enrichment of the genes differentially expressed in OE5 relative to all remaining cells of the same cell type; bars show -log10(P).

**Supplementary Figure S23.** Pathway enrichment for region R1 (reticulocytes). Metascape enrichment of the genes differentially expressed in R1 relative to all remaining cells of the same cell type; bars show -log10(P).

**Supplementary Figure S24.** Pathway enrichment for region R2 (reticulocytes). Metascape enrichment of the genes differentially expressed in R2 relative to all remaining cells of the same cell type; bars show -log10(P).

**Supplementary Figure S25.** Pathway enrichment for region R3 (reticulocytes). Metascape enrichment of the genes differentially expressed in R3 relative to all remaining cells of the same cell type; bars show -log10(P).

### Supplementary Figure S26. InferCNV score distributions and derivation of the score threshold.

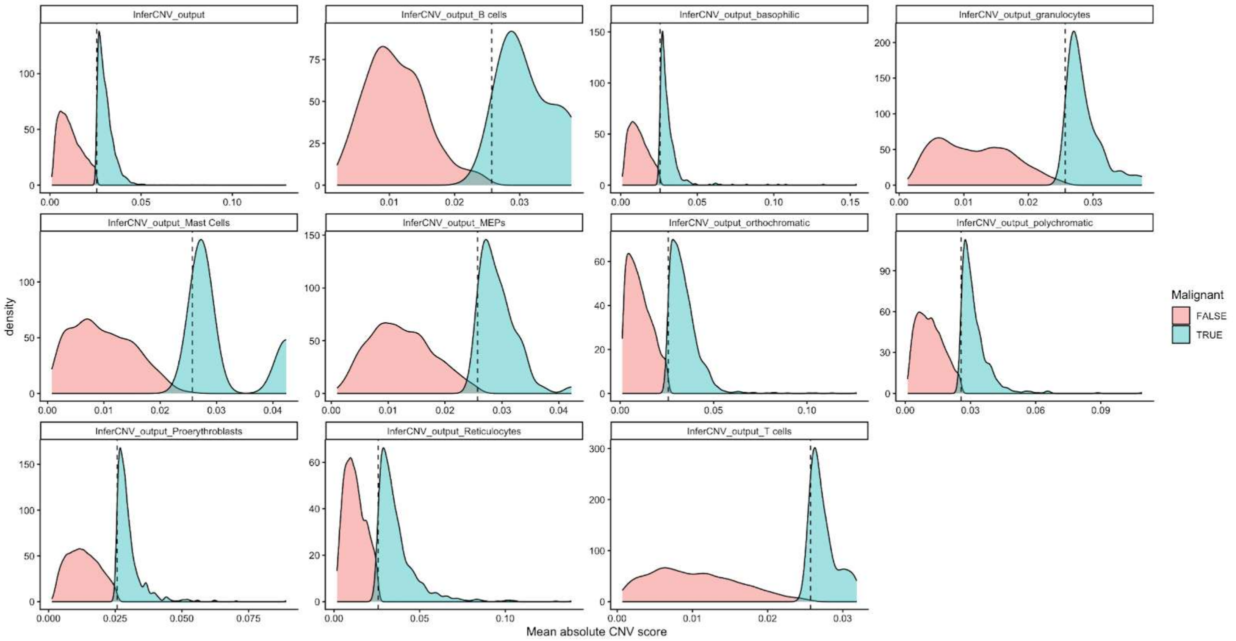


Density of the mean absolute inferCNV score for the whole object and for each annotated cell type, split into cells above (TRUE) and below (FALSE) the threshold. The dashed line marks the threshold of 0.02570775, defined as the maximum of the reference-cell median plus three median absolute deviations and the reference-cell 99th percentile. Reference populations were B cells, T cells, granulocytes and mast cells.

### Supplementary Tables

**Supplementary Table S1. Bulk differential expression results (GSE26049).** 324 differentially expressed genes (89 up, 235 down in PV) from 41 PV and 21 control samples; limma v3.54.0, $|\log2FC| > 1.0$, adjusted $P < 0.05$.

**Supplementary Table S2. Disease-effect and treatment-effect gene lists per cell type.** Complete disease-effect and treatment-effect gene symbol lists for all ten annotated cell types. The basophilic erythroblast disease-effect list contains the full 53 genes, replacing the run-on sentence previously carried in the manuscript Results.

**Supplementary Table S3. Pseudotime trajectory regions.** All 19 trajectory regions (BE1–BE6, PE1–PE5, OE1–OE5, R1–R3) with dominant condition and enrichment theme. T-cell regions are excluded, matching the manuscript.

**Supplementary Table S4. Differential expression statistics for each trajectory region.** Per-region differential expression statistics underlying the enrichment figures S7–S25.

**Supplementary Table S5. Genes correlated with inferCNV score.** The nine genes at the 16p13.3 alpha-globin locus most strongly correlated with the per-cell CNV score (mean absolute Spearman correlation > 0.13).

**Supplementary Table S6. Basophilic erythroblast versus proerythroblast differential expression.** Genes differentially expressed at the proerythroblast to basophilic erythroblast transition, including the coordinated decline of KIT, STAT5A, CDK6, CASP3 and SMARCC1.

**Supplementary Table S7. Drug repurposing candidates.** Ruxolitinib plus interferon alpha, navitoclax, and SMARCC1 as a proposed but currently undrugged target, each with mechanism and trial evidence.
